# CRISPR/Cas9-engineered inducible gametocyte producer lines as a novel tool for basic and applied research on *Plasmodium falciparum* malaria transmission stages

**DOI:** 10.1101/2020.10.05.326868

**Authors:** Sylwia D. Boltryk, Armin Passecker, Arne Alder, Marga van de Vegte-Bolmer, Robert W. Sauerwein, Nicolas M. B. Brancucci, Hans-Peter Beck, Tim-Wolf Gilberger, Till S. Voss

## Abstract

The malaria parasite *Plasmodium falciparum* replicates inside erythrocytes in the blood of infected humans. During each replication cycle, a small proportion of parasites commits to sexual development and differentiates into gametocytes, which are essential for parasite transmission to other human hosts via the mosquito vector. Detailed molecular investigation of gametocyte biology and transmission has been hampered by difficulties in generating large numbers of these highly specialized cells. Here, we engineered marker-free *P. falciparum* inducible gametocyte producer (iGP) lines for the routine mass production of synchronous gametocytes. Through targeted overexpression of the sexual commitment factor GDV1, iGP lines consistently achieve sexual commitment rates of 75% and produce gametocytes that are infectious to mosquitoes. Subsequent tagging of a nucleoporin allowed us to visualize marked nuclear transformations during gametocytogenesis and demonstrates that further genetic engineering of iGP lines is an invaluable tool for the targeted exploration of gametocyte biology. We believe the iGP approach developed here opens up unprecedented opportunities that will expedite future basic and applied research on *P. falciparum* transmission stages.

## Introduction

Malaria is a vector-borne infectious disease caused by protozoan parasites of the genus *Plasmodium*. Infections with *P. falciparum* are responsible for the vast majority of all malaria-related morbidity and mortality in humans. Over the last two decades concerted intervention efforts targeting both the insect vector and the parasite led to a remarkable decline in malaria cases worldwide ^1^. However, progress has come to a standstill in the past few years and in 2018 malaria was still accountable for 228 million clinical cases and 405’000 deaths, primarily in sub-Saharan Africa ^2^. To further reduce the spread of malaria, intervention strategies will not only have to overcome the widespread resistance of mosquitoes and parasites to insecticides and first-line antimalarial drugs, respectively, but will also have to include efficient tools that interrupt parasite transmission from the human host to the mosquito vector ^3,4^.

People get infected with malaria parasites when *P. falciparum*-infested female *Anopheles* mosquitoes inject sporozoites into their skin. After reaching the liver via the bloodstream, sporozoites multiply within hepatocytes to release thousands of merozoites into circulation. Merozoites invade red blood cells (RBCs) and develop through the ring and trophozoite stage into a multi-nucleated schizont. After daughter cell formation, up to 32 merozoites egress from each infected RBC (iRBC) to invade and replicate inside new RBCs. Consecutive rounds of these intra-erythrocytic developmental cycles (IDCs) are responsible for all disease symptoms and chronic infection. Importantly, however, during each replication cycle a small proportion of schizonts produce sexually committed ring stage progeny that differentiate into gametocytes ^5^. When taken up by a mosquito, terminally differentiated gametocytes egress from the iRBC and develop into gametes. After fertilization, the zygote transforms into an ookinete that traverses the midgut epithelium to initiate sporogony, which ultimately renders the mosquito infectious to other humans. Hence, as the only forms of the parasite able to infect mosquitoes, gametocytes are highly specialized cells that secure human-to-human malaria transmission ^6^.

In *P. falciparum*, the process of sexual differentiation takes 10-12 days during which the parasite undergoes drastic changes in cellular morphology. Sexual ring stages (day 1) develop into spherical stage I gametocytes (day 2) that continuously elongate into lemon-shaped stage II (day 4), D-shaped stage III (day 6) and spindle-shaped stage IV cells (day 8), before falciform mature stage V gametocytes are formed (day 10+) ^7,8^. These morphological transitions are linked to the gradual expansion of the inner membrane complex, an endomembrane system underlying the parasite plasma membrane, and microtubule and actin cytoskeleton networks underneath that are disassembled again at the stage IV to V transition ^9-13^. Stage I to IV gametocytes are sequestered away from circulation, primarily in the parenchyma of the bone marrow and spleen ^6,14^. The mechanisms involved in gametocyte homing to and sequestration in these extravascular niches are poorly understood. However, the high rigidity of stage I to IV gametocyte-infected RBCs ^9,15,16^, established through parasite-induced alterations of the RBC membrane and underlying cytoskeletal networks ^15,17-19^, appears to play a primary role in gametocyte retention. Reversal of these modifications at the stage IV to V transition confers increased deformability ^9,15-19^, which likely allows for the release of stage V gametocytes into the bloodstream ^6,20^. Once in circulation, stage V gametocytes remain competent for transmission to the mosquito for days/weeks ^21^. The enormous transmission reservoir represented by the hundreds of millions of infected people in endemic areas, and the failure of almost all currently licensed antimalarial drugs except primaquine to efficiently kill mature gametocytes, pose major obstacles to malaria control and elimination efforts ^4,22,23^. Furthermore, the development of transmission-blocking drugs and vaccines is compromised by our poor understanding of the molecular mechanisms underlying essential gametocyte biology and the lack of reliable and readily applicable experimental tools.

Parasites commit to gametocytogenesis during the IDC preceding gametocyte differentiation. This finding was made based on experiments showing that all ring stage descendants derived from a single schizont have the same fate; they either all undergo another round of intracellular replication or they all differentiate into either female or male gametocytes ^24,25^. In addition to this ‘next cycle sexual conversion (NCC)’ process, recent studies reported ‘same cycle sexual conversion (SCC)’ where ring stages directly commit to sexual development ^26,27^. Irrespective of the NCC or SCC routes, sexual conversion is triggered by an epigenetic switch that activates expression of the master transcription factor AP2-G ^28-31^. In asexual parasites, heterochromatin-dependent silencing of *ap2-g* prevents AP2-G expression ^28,30-32^. In a small subset of trophozoites (NCC) or ring stages (SCC), however, the *ap2-g* locus gets activated by molecular mechanisms that are still largely unknown ^27-31,33-35^. Gametocyte development 1 (GDV1), a nuclear protein essential for gametocytogenesis in *P. falciparum* ^36^, plays a key role in the NCC process ^34^. GDV1 is specifically expressed in sexually committed parasites, where it displaces heterochromatin protein 1 (HP1) from the *pfap2-g* locus thereby licensing PfAP2-G expression ^34^. Consistent with the epigenetic control mechanisms regulating *pfap2-g* expression, sexual commitment rates vary in response to various environmental conditions ^5^. In particular, depletion of the host serum lipid lysophosphatidylcholine (LysoPC) triggers sexual commitment and this response is channeled via induction of GDV1 and PfAP2-G expression ^34,35^. Once expressed, PfAP2-G initiates a specific transcriptional programme that drives sexual conversion and subsequent gametocyte differentiation ^5^. While the PfAP2-G-dependent transcriptional changes in sexually committed schizonts are minor, a more pronounced gene expression signature emerges in the sexually committed ring stage progeny, where several dozen genes are specifically induced or repressed compared to asexual ring stages ^28,30,31,33,34,37,38^. Most of these upregulated genes are directly targeted by PfAP2-G ^33^ and encode proteins implicated in iRBC remodeling, while some others have expected roles in regulating gene expression during gametocyte maturation ^28,30,31,33,34,37^.

Regarding the complex process of sexual differentiation, mRNA and protein expression profiling studies conducted mainly on late stage gametocytes ^39-43^, but also on early stages ^42,43^ or across gametocyte maturation ^44,45^, identified hundreds of genes and proteins specifically up- or downregulated compared to asexual blood stages, or between female and male gametocytes ^46-49^. Stage I/II gametocytes express exported proteins involved in sexual stage-specific iRBC remodeling ^42^. Enzymes of the oxidative tricarboxylic acid (TCA) cycle are upregulated in gametocytes ^39,44,45^, consistent with their increased dependence on the TCA cycle for energy production ^50^. Factors involved in protein synthesis, mitochondrial function and translational repression are upregulated in females ^46-49^, in line with the increased abundance of ribosomes ^12^, enlarged mitochondrion ^12,51^ and large number of translationally repressed transcripts in these stages ^46^. Proteins linked to DNA replication and axoneme formation are enriched in males to prepare for the three rapid rounds of genome duplication and generation of eight motile microgametes during male gametogenesis ^46-48^. Gametocytes also show increased expression of transcription factors and chromatin modifiers ^28,30,33,39,44,45^, expanded subtelomeric heterochromatin domains ^32,52^, distinct histone post-translational modification profiles ^53^ and altered three-dimensional chromosome organisation ^52^. Furthermore, earlier studies investigating gametocyte morphology at the ultrastructural level described an intriguing nuclear dimorphism between female and male gametocytes ^12,54^. These nuclear processes and features have not been further explored in any great detail but are likely linked to the control of gametocyte and sex-specific differentiation. These and many additional studies ^5,55^ provided invaluable insight into gametocyte- and sex-specific biology. However, a functional and mechanistic understanding of the underlying molecular and cellular processes is often lacking since only a relatively small number of genes has been investigated by reverse genetics approaches in *P. falciparum* gametocytes.

Our limited knowledge about gametocyte biology is primarily due to the difficulty in generating large numbers of synchronous gametocyte stages for experimental studies: (i) *in vitro* cultured *P. falciparum* parasites show low sexual commitment rates (<10%) (reflected as the proportion of sexually committed ring stages among all ring stage parasites) ^5,6^; (ii) gametocytes are non-proliferative cells and thus rapidly overgrown by asexual parasites; and (iii) sexual commitment occurs during each consecutive IDC, which results in asynchronous gametocyte populations. Currently used approaches to increase sexual commitment rates for *in vitro* gametocyte production are based on exposing parasite cultures to poorly defined stress conditions imposed for instance by high parasitaemia and/or nutrient starvation (spent/conditioned medium) ^8,56,57^. Several different protocols relying on this strategy exist and reach reported sexual commitment rates of 10-30% ^58-64^. However, most of these protocols use cumbersome experimental workflows, rely on large culture volumes, are expensive, are difficult to reproduce and/or produce asynchronous gametocyte populations. The recently developed approach employing LysoPC- or choline-depleted minimal fatty acid medium^35^ achieves sexual commitment rates in the range of 15-60%, but growth under these nutrient-restricted conditions leads to lower numbers of sexually committed progeny produced per schizont ^34,35,65,66^.

Here, we used CRISPR/Cas9-based genome editing to engineer transgenic *P. falciparum* lines suitable for the routine mass production of synchronous gametocytes. Through the targeted induction of GDV1 expression these lines consistently achieve sexual conversion rates of 75% and generate synchronous gametocyte populations at high yield across all stages of gametocytogenesis. Furthermore, we demonstrate that these gametocytes undergo sexual reproduction and oocyst formation in female *Anopheles* mosquitoes. Lastly, by generating iGP parasites expressing an mScarlet-tagged version of the nuclear pore protein NUP313 we show that further genetic engineering of iGP lines is straightforward and enables the generation of large numbers of gametocyte mutants for the targeted investigation of *P. falciparum* transmission stage biology.

## Results

### CRISPR/Cas9-based engineering of an inducible gametocyte producer line in the 3D7 reference strain

Using the FKBP destabilization domain (DD) system for controllable protein expression ^67^, we previously observed that conditional overexpression of plasmid-encoded GDV1 increases sexual conversion rates ^34^. In this study, we aimed to exploit this finding in order to generate stable and marker-free inducible gametocyte producer (iGP) lines as tools to expedite research on *P. falciparum* gametocyte biology and transmission. As a proof of concept, we inserted a single GDV1-GFP-DD expression cassette into the dispensable *cg6* (*glp3*) locus (PF3D7_0709200) ^68^ in the 3D7 reference strain (Fig. 1a). To this end, we co-transfected the pHF_gC-*cg6* CRISPR/Cas9 plasmid (containing expression cassettes for the positive-negative selection marker human dihydrofolate reductase fused to yeast cytosine deaminase/uridyl phosphoribosyl transferase (hDHFR-yFCU), SpCas9 and the single guide RNA) and the pD_*cg6_cam-gdv1-gfp-dd* donor plasmid into 3D7 wild type parasites (Fig. S1). Transgenic 3D7/iGP parasites were readily selected on WR99210 and PCRs on genomic DNA (gDNA) confirmed complete disruption of the *cg6* locus through insertion of the *gdv1-gfp-dd* donor assembly. At this stage, both transfected plasmids were still detectable in the population and the majority of parasites carried integrated donor plasmid concatamers (Fig. S1), which has also been reported in other studies ^69,70^. Treatment with 5-fluorocytosine (5-FC) successfully depleted parasites expressing the hDHFR-yFCU marker and enriched for parasites carrying a single GDV1-GFP-DD expression cassette integrated into the *cg6* locus (Fig. S1). This population was then used to obtain three 3D7/iGP clonal lines by limiting dilution cloning (Fig. S1). PCRs on gDNA demonstrated that clones 3D7/iGP_B9 and 3D7/iGP_F10 are clean plasmid- and marker-free parasites carrying a single integrated GDV1-GFP-DD expression cassette. Clone 3D7/iGP_D9 is also marker-free but still contains a donor plasmid concatamer in the *cg6* locus (Fig. S1).

**Figure 1.**
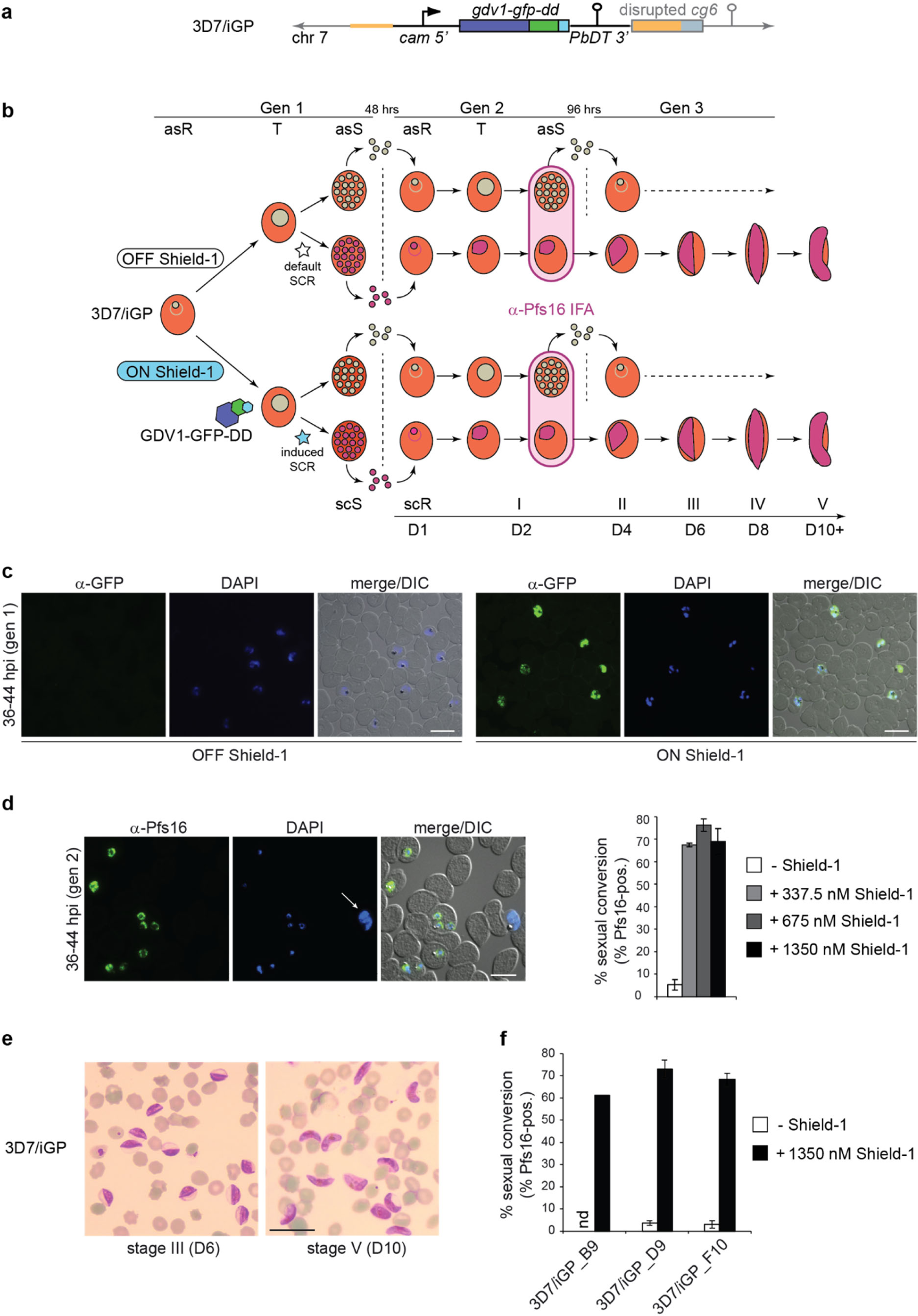
Description of the inducible gametocyte producer line 3D7/iGP. **a** Schematic map of the disrupted *cg6* (*glp3*) locus (PF3D7_0709200) (grey) carrying a single inducible GDV1-GFP-DD expression cassette (black) in 3D7/iGP parasites. The 5’ and 3’ homology regions used for CRISPR/Cas9-based transgene insertion are positioned in the *cg6* upstream and coding sequence, respectively (orange) (see also Supplementary Fig. 1). **b** Schematic representation of the *in vitro* culture protocol used to quantify sexual commitment rates in 3D7/iGP parasites. Synchronous 3D7/iGP ring stage parasites are split at 0-16 hpi and Shield-1 is added to one half of the population to trigger GDV1-GFP-DD expression (generation 1). Sexual commitment rates (SCR) are quantified by determining the proportion of early stage I gametocytes in the total iRBC progeny 36-44 hpi (generation 2; day 2 of gametocytogenesis) by α-Pfs16 IFAs combined with DAPI staining. Asexual parasites are depicted in grey, sexually committed parasites and gametocytes are depicted in purple. asS/scS, asexual/sexually committed schizont; asR/scR, asexual/sexually committed ring stage; T, trophozoite; I-V, gametocyte stages I-V; D1-D10, days 1-10 of gametocyte maturation. Gen1/2, generation 1/2. **c** Representative α-GFP IFA images illustrating the Shield-1-dependent induction of GDV1-GFP-DD expression in schizonts of the 3D7/iGP mother line (generation 1). Nuclei were stained with DAPI. DIC, differential interference contrast. Scale bar, 10 µm. **d** Representative α-Pfs16 IFA images illustrating the high proportion of early gametocytes in the progeny of the Shield-1-treated 3D7/iGP mother line. The white arrow highlights a Pfs16-negative schizont. Nuclei were stained with DAPI. DIC, differential interference contrast. Scale bar, 10 µm (left panel). Proportion of Pfs16-positive iRBCs (sexual conversion rate) in the progeny of the 3D7/iGP mother line treated with three different Shield-1 concentrations and the untreated control. Values are the mean (± s.d.) of three independent biological replicates (two replicates for populations treated with 337.5 nM Shield-1) (right panel) (>200 DAPI-positive cells counted per experiment). **e** Representative images of Giemsa-stained gametocyte cultures obtained after inducing sexual commitment in the 3D7/iGP mother line, acquired on day 6 (stage III gametocytes) and day 10 (stage V gametocytes) of gametocytogenesis. Scale bar, 20 µm. **f** Proportion of Pfs16-positive iRBCs (sexual conversion rate) in the progeny of untreated (control) or Shield-1-treated 3D7/iGP clones. Values are the mean (± s.d.) of three independent biological replicates (only one experiment performed for clone B9) (>300 DAPI-positive cells counted per experiment).

To test if Shield-1 induces GDV1-GFP-DD overexpression and sexual conversion, we split ring stage parasites of the 5-FC-treated 3D7/iGP line at 8-16 hours post-invasion (hpi) (generation 1) into two equal populations and added 1350 nM Shield-1 to one of them (Fig. 1b). Indirect immunofluorescence assays (IFA) on schizont stage parasites (40-48 hpi, generation 1) showed that most parasites in the Shield-1-treated population expressed GDV1-GFP-DD, in contrast to parasites cultured in absence of Shield-1 where GDV1-GFP-DD expression was hardly detectable (Fig. 1c). IFAs probing for expression of the gametocyte marker Pfs16 ^71^ in the early stage I gametocyte progeny (36-44 hpi, generation 2; day 2 of gametocytogenesis) revealed mean sexual conversion rates of 5.2% (± 1.9 s.d.) and 69.0% (± 4.5 s.d.) in parasites cultured in absence and presence of Shield-1, respectively (Fig. 1d). Similarly high sexual conversion rates were achieved when Shield-1 was used at 675 nM or 337.5 nM (Fig. 1d). Furthermore, all three 3D7/iGP clones showed high sexual conversion rates comparable to those achieved with the 3D7/iGP mother line (Fig. 1e). To test if Shield-1-induced 3D7/iGP early stage gametocytes complete sexual differentiation, the sexual ring stage progeny was exposed to 50 mM N-acetylglucosamine (GlcNAc) for six consecutive days (days 1-6 of gametocytogenesis) to eliminate asexual parasites ^58,72^ and was thereafter maintained under routine culture conditions. Inspection of Giemsa-stained blood smears prepared on days 6 and 10 of gametocyte maturation revealed pure and synchronous stage III and V gametocyte populations, respectively (Fig. 1f). Hence, by integrating a conditional GDV1-GFP-DD overexpression cassette into the 3D7 genome we obtained a stable 3D7/iGP line and marker-free clones suitable for the production of synchronous gametocyte cultures at high yield.

### CRISPR/Cas9-based engineering of inducible gametocyte producer lines in the transmissible NF54 strain

We next wanted to transfer the iGP system to NF54 parasites, the strain most widely used for the study of gametocyte biology and mosquito infection ^73^. Unexpectedly, however, multiple attempts to insert the GDV1-GFP-DD expression cassette into the *cg6* locus in NF54 parasites failed as we never obtained viable parasites after drug selection of transfected populations. While the reason for this failure is unknown, we speculated that the DD-dependent degradation of GDV1-GFP-DD might be less efficient in the NF54 strain such that transfected parasites would commit to sexual differentiation at a high rate even in absence of Shield-1 and therefore fail to proliferate. Based on this speculation we decided to employ the *glmS* riboswitch approach that allows regulating mRNA levels in *P. falciparum* ^74^. When located in the untranslated region of mRNA, the *glmS* ribozyme triggers transcript degradation in presence of glucosamine (GlcN). We generated two modified donor plasmids by (1) adding a *glmS* ribozyme element downstream of the *gdv1-gfp-dd* sequence (pD_*cg6_cam-gdv1-gfp-dd-glmS*); and (2) by removing the *dd* sequence and adding a *glmS* ribozyme element instead (pD_*cg6_cam-gdv1-gfp-glmS*) (Fig. S2). Co-transfection of the pBF_gC-*cg6* CRISPR/Cas9 plasmid (which carries the blasticidin deaminase (*bsd*) gene as a positive selection marker instead of h*dhfr*) (Fig. S2) with either of the donor plasmids allowed us to successfully select for transgenic NF54/iGP populations in both instances. PCRs on gDNA of the NF54/iGP1 line revealed complete disruption of the *cg6* locus and insertion of a single GDV1-GFP-DD-*glmS* expression cassette (Fig. S2). The CRISPR/Cas9 plasmid and integrated donor plasmid concatamers were also detectable in the population. While treatment with 5-FC eliminated parasites harbouring the pBF_gC-*cg6* plasmid but not those carrying integrated donor plasmid concatamers, five NF54/iGP1 clones obtained from this population were all plasmid- and marker-free and carried a single GDV1-GFP-DD-*glmS* expression module in the *cg6* locus (Fig. S2). The NF54/iGP2 population was plasmid- and marker-free showing complete disruption of the *cg6* locus through insertion of a single GDV1-GFP-*glmS* expression cassette, and the same result was obtained for four NF54/iGP2 clones (Fig. S2).

Induction of GDV1-GFP-DD expression in the NF54/iGP1 line by the simultaneous removal of GlcN and addition of Shield-1 in generation 1 produced progeny consisting of 74.8% (±4.6 s.d.) Pfs16-positive early stage I gametocytes (32-40 hpi in generation 2) compared to only 8.0% (±2.6 s.d.) in the control population (Figs. 2a and 2b). The same induction protocol applied to three NF54/iGP1 clones resulted in somewhat lower but still remarkable sexual conversion rates of up to 60% (Fig. 2b). Likewise, induction of GDV1-GFP expression in the NF54/iGP2 line via removal of GlcN delivered progeny consisting of 74.1% (±5.6 s.d.) early stage I gametocytes compared to only 5.7% (±1.3 s.d.), and slightly lower sexual conversion rates were observed for three NF54/iGP2 clones (Figs. 2c and 2d).

**Figure 2.**
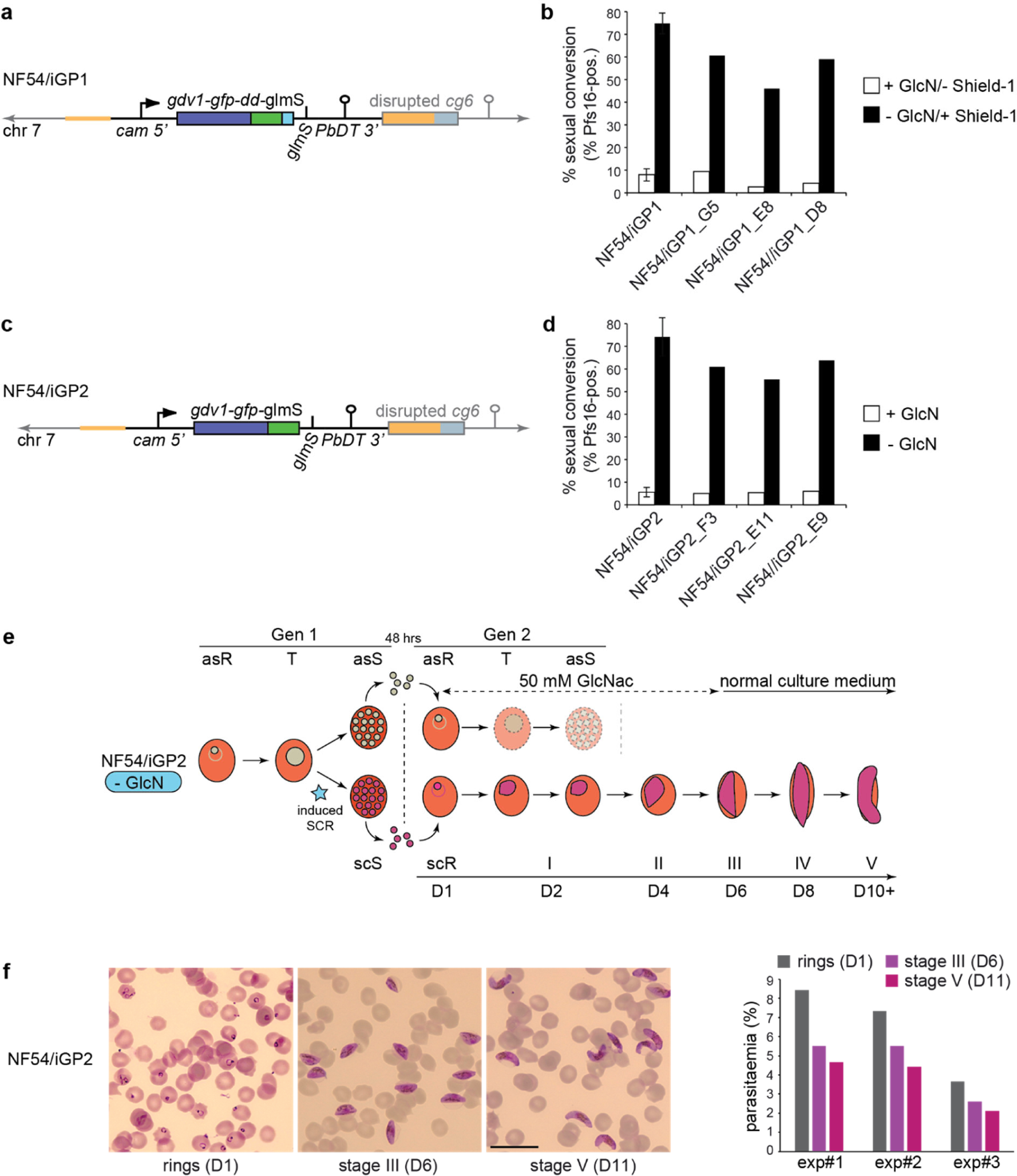
Description of the inducible gametocyte producer lines NF54/iGP1 and NF54/iGP2. **a** Schematic map of the disrupted *cg6* (*glp3*) locus (PF3D7_0709200) (grey) carrying a single inducible GDV1-GFP-DD-glmS expression cassette (black) in NF54/iGP1 parasites. The 5’ and 3’ homology regions used for CRISPR/Cas9-based transgene insertion are positioned in the *cg6* upstream and coding sequence, respectively (orange) (see also Supplementary Fig. 2). **b** Proportion of Pfs16-positive iRBCs (sexual conversion rate) in the progeny of the NF54/iGP1 mother line and three clones cultured in the presence of 2.5 mM GlcN/absence of Shield-1 (control) or in the absence of GlcN/presence of 1350 nM Shield-1. Values represent the mean (± s.d.) of three independent biological replicates (NF54/iGP1 mother line) or the result of a single experiment (NF54/iGP1 clones) (>300 DAPI-positive cells counted per experiment). **c** Schematic map of the disrupted *cg6* (*glp3*) locus (PF3D7_0709200) (grey) carrying a single inducible GDV1-GFP-glmS expression cassette (black) in NF54/iGP2 parasites (see also Supplementary Fig. 2). **d** Proportion of Pfs16-positive iRBCs (sexual conversion rate) in the progeny of the NF54/iGP2 mother line and three clones cultured in the presence of 2.5 mM GlcN (control) or in the absence of GlcN. Values represent the mean (± s.d.) of three independent biological replicates (NF54/iGP2 mother line) or the result of a single experiment (NF54/iGP2 clones) (>300 DAPI-positive cells counted per experiment). **e** Schematic representation of the *in vitro* culture protocol used to obtain pure NF54/iGP2 stage V gametocyte populations. GlcN is removed from the culture medium of synchronous NF54/iGP2 ring stage parasites at 0-16 hpi to trigger expression of GDV1-GFP (generation 1). After schizont rupture and merozoite invasion gametocyte maturation is allowed to proceed for >10 days. 50 mM GlcNAc is added to the culture medium for the first six days of gametocytogenesis to eliminate asexual parasites. Note: for NF54/iGP1 parasites, GlcN is removed and Shield-1 added to the culture medium of synchronous ring stage parasites (0-16 hpi) to trigger expression of GDV1-GFP-DD (generation 1). Asexual parasites are depicted in grey, sexually committed parasites and gametocytes are depicted in purple. asS/scS, asexual/sexually committed schizont; asR/scR, asexual/sexually committed ring stage; T, trophozoite; I-V, gametocyte stages I-V; D1-D10, days 1-10 of gametocyte maturation. Gen1/2, generation 1/2. **f** Representative images of Giemsa-stained gametocyte cultures obtained after inducing sexual commitment in the NF54/iGP2 mother line, acquired on day 1 (asexual/sexually committed ring stages), day 6 (stage III gametocytes) and day 11 (stage V gametocytes) of gametocytogenesis. Scale bar, 20 µm. Parasitaemia as determined by visual inspection of Giemsa-stained thin blood smears prepared on day 1, day 6 and day 11 of gametocytogenesis from three independent gametocyte induction experiments starting at different parasitaemias are shown on the right.

To monitor gametocyte maturation, the ring stage progeny of induced NF54/iGP1 and NF54/iGP2 parasites were cultured in medium containing 50 mM GlcNAc for six days and then maintained under routine culture conditions until day 12 of gametocyte development (Fig. 2e). Visual inspection of Giemsa-stained blood smears prepared daily from day 4 (stage II) onwards revealed that these gametocytes differentiated in a highly synchronous manner into stage V gametocytes within 10-12 days (Fig. 2f and Fig. S3). Notably, due to the high sexual conversion rates achieved, pure gametocyte populations at 4-5% gametocytaemia were routinely obtained (Fig. 2f). In summary, by applying the *glmS* riboswitch system for the conditional activation of GDV1 overexpression, we were able to engineer two independent marker-free NF54/iGP lines that allow for the routine production of large numbers of pure and synchronous NF54 gametocytes across all stages of sexual differentiation using a most simple induction protocol.

### NF54/iGP1 and NF54/iGP2 gametocytes are infectious to mosquitoes

To test if NF54/iGP1 and NF54/iGP2 gametocytes retained their capacity to infect mosquitoes we fed stage V gametocyte preparations to female *Anopheles stephensi* mosquitoes using standard membrane feeding assays (SMFAs) ^73,75^. To this end, sexual conversion was induced in clones NF54/iGP1_D8 and NF54/iGP2_E9 by addition of Shield-1/removal of GlcN or removal of GlcN, respectively, and the ring stage progeny (day 1 of gametocytogenesis) was maintained in culture for 14 days with daily medium changes to ensure complete differentiation into mature stage V gametocytes. For each of the two populations, separate SMFAs were performed on days 10, 13 and 14 of gametocytogenesis. Eight days later, 20 mosquitoes each were dissected and midgut oocysts counted by microscopy. As shown in Fig. 3, the infection rates achieved with both clones were high for each feed, with 80-100% and 90-100% of mosquitoes successfully infected by NF54/iGP1_D8 and NF54/iGP2_E9 stage V gametocytes, respectively, on each of the three days of feeding. Mosquitoes infected with NF54/iGP2_E9 gametocytes consistently developed higher numbers of oocysts per mosquito compared to those fed with NF54/iGP1_D8 gametocytes. NF54/iGP1_D8 gametocytes scored the highest oocyst loads when fed on day 13 of gametocytogenesis (median=18 oocysts/mosquito), whereas NF54/iGP2_E9 gametocytes were most infectious on days 13 (median=32 oocysts/mosquito) and 14 (median=35 oocysts/mosquito) (Fig. 3). Importantly, the infection rates and oocyst loads achieved with NF54/iGP1_D8 and NF54/iGP2_E9 stage V gametocytes are comparable to those obtained with NF54 wild type gametocytes^76^

**Figure 3.**
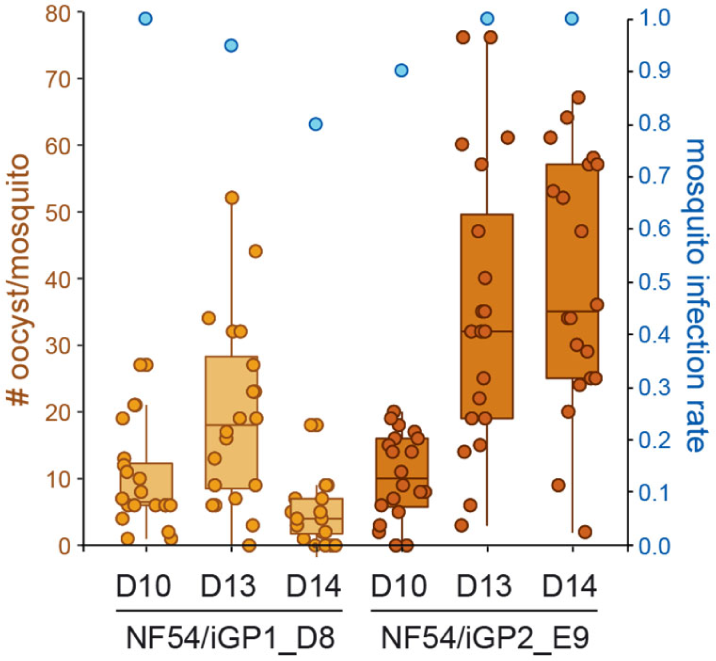
NF54/iGP1 and NF54/iGP2 gametocytes infect mosquitoes. Stage V gametocytes derived from clones NF54/iGP1_D8 (beige) and NF54/iGP2_E9 (brown) were fed to female *Anopheles stephensi* mosquitoes on day 10, 13 and 14 of gametocytogenesis (D10, D13, D14). The number of oocysts detected in each of the 20 mosquitoes dissected per feeding experiment are shown as beige and brown circles, and the median number of oocysts detected among the 20 mosquitoes per feeding experiment (horizontal line) and upper and lower quartile ranges are represented by box plots (left y-axis). The proportion of infected mosquitoes obtained per feeding experiments is indicated by blue circles (right y-axis).

### Further genetic manipulation of iGP parasites for the targeted investigation of gametocyte biology

To explore iGP parasites as a tool to study specific aspects of gametocyte biology, we modified NF54/iGP2 parasites again using a second CRISPR/Cas9 gene editing step. Based on observations made by thin-section transmission electron microscopy (TEM) regarding the increased size and altered shape of male gametocyte nuclei ^12,54^, we were interested in visualising the outer confines of the parasite nucleus throughout gametocytogenesis by fluorescence microscopy. Since *P. falciparum* lacks nuclear lamin, and potential homologs of known nuclear envelope transmembrane proteins have not yet been described, we tagged a protein of the nuclear pore complex (NPC) as a surrogate marker for the nuclear membrane. Nuclear pores are large macromolecular complexes consisting of approximately 30 different nucleoporins (NUPs) (each present in multiple copies) that are embedded in the nuclear envelope (Field and Rout, 2019. NPCs act as essential gateways for molecular transport into and out of the nucleus and play additional crucial roles in the regulation of gene expression and genome organization ^77,78^. In malaria parasites, nuclear pores have hardly been studied and only six NUPs have so far been identified ^79-83^, of which four have been localized in *P. falciparum* asexual blood stage parasites [PfNUP116 (PF3D7_1473700) ^80^, PfNUP221/PfNUP100 (PF3D7_0905100) ^83^, PfNUP313 (PF3D7_1446500) ^79^, PfSEC13 (PF3D7_1230700) ^82^].

To visualize NPC distribution in gametocytes we engineered NF54/iGP2 parasites expressing mScarlet-tagged NUP313 (NF54/iGP2_NUP313-mSc). To achieve this, we co-transfected the pBF_gC-*nup313* CRISPR/Cas9 plasmid and the pD_*nup313-mScarlet* donor plasmid into the marker-free NF54/iGP2 line (Fig. S4). Transgenic parasites were successfully selected on BSD-S-HCl and PCRs on gDNA confirmed correct editing of the *nup313* gene and absence of parasites carrying the wild type locus (Fig. S4). Validation of the NF54/iGP2_NUP313-mSc line by live cell fluorescence imaging verified the expected localization of nuclear pores across the different IDC stages as determined in previous studies ^80,84,85^ (Fig. 4a). NUP313-mScarlet localized to a single region adjacent to the DAPI-stained area in merozoites and early ring stages. The increased number and even distribution of NPCs within the nuclear envelope in trophozoites and early schizonts was reflected by a circular perinuclear pattern of NUP313-mScarlet foci directly adjoining the genetic material. In late schizonts, the number of NUP313-mScarlet signals decreased again to one or two per nucleus. To assess NUP313-mScarlet localization in gametocytes we induced sexual commitment in NF54/iGP2_NUP313-mSc parasites according to the protocol outlined in Fig. 2e and performed live cell fluorescence imaging for all five gametocyte stages. As shown in Fig. 4b, NUP313-mScarlet signals were abundant and surrounded the often elongated area of nuclear DNA in a dot-like fashion in all five gametocyte stages, similar to the localization pattern observed in trophozoites (Fig. 4a). However, in stage II to V gametocytes the NPC signals frequently stretched away from bulk chromatin, suggestive of rounded expansions (pink arrowheads) and narrow lateral extensions (yellow arrowheads) of the nuclear envelope that sometimes reached close to the cellular poles (Figs. 4b and 5, Fig. S5). Primarily in late stage gametocytes we also observed nuclei with two clearly separate NUP313-delineated regions, of which either both (white arrowheads) or only one (blue arrowhead) contained genetic material detectable by Hoechst-staining (Fig. 5). In summary, our findings show that gametocyte nuclei contain a similar or even higher density of NPCs compared to trophozoites, undergo profound changes in nuclear shape and size and often contain regions of low chromatin density.

**Figure 4.**
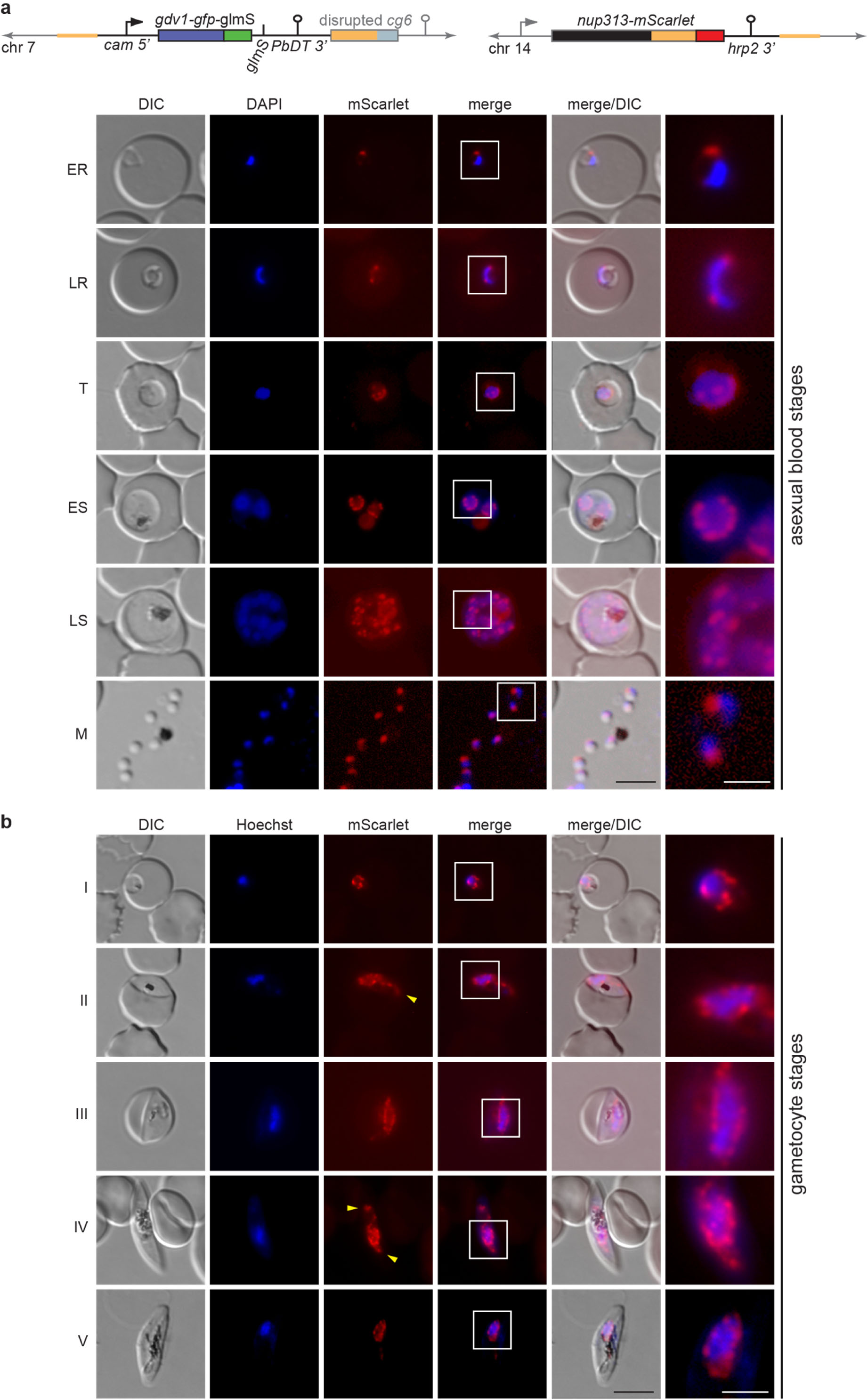
Visualisation of nuclear pore distribution in NF54/iGP2_NUP313-mSc asexual blood stage parasites and gametocytes. **a** Schematic maps of the disrupted *cg6* (*glp3*) locus (PF3D7_0709200) (grey) carrying a single inducible GDV1-GFP-glmS expression cassette and the tagged *nup313* locus in double-transgenic NF54/iGP2_NUP313-mSc line are shown on top. The 5’ and 3’ homology regions used for CRISPR/Cas9-based gene editing are indicated in orange (see also Figs. S2 and S4). Representative live cell fluorescence microscopy images showing the localization of NUP313-mScarlet (red) in asexual blood stage parasites. ER/LR, early/late ring stage; T, trophozoite; ES/LS, early/late schizont; M, merozoite. DIC, differential interference contrast. Nuclei were stained with DAPI (blue). Scale bar, 5 µm. White frames refer to the magnified view presented in the rightmost images (scale bar, 2 µm). **b** Representative live cell fluorescence microscopy images showing the localization of NUP313-mScarlet (red) in stage I to V gametocytes. Lateral extensions of the nucleus away from Hoechst-stained bulk chromatin are highlighted by yellow arrowheads. I-V, stage I to V gametocytes. DIC, differential interference contrast. Nuclei were stained with Hoechst (blue). Scale bar, 5 µm. White frames refer to the magnified view presented in the rightmost images (scale bar, 2 µm).

**Figure 5.**
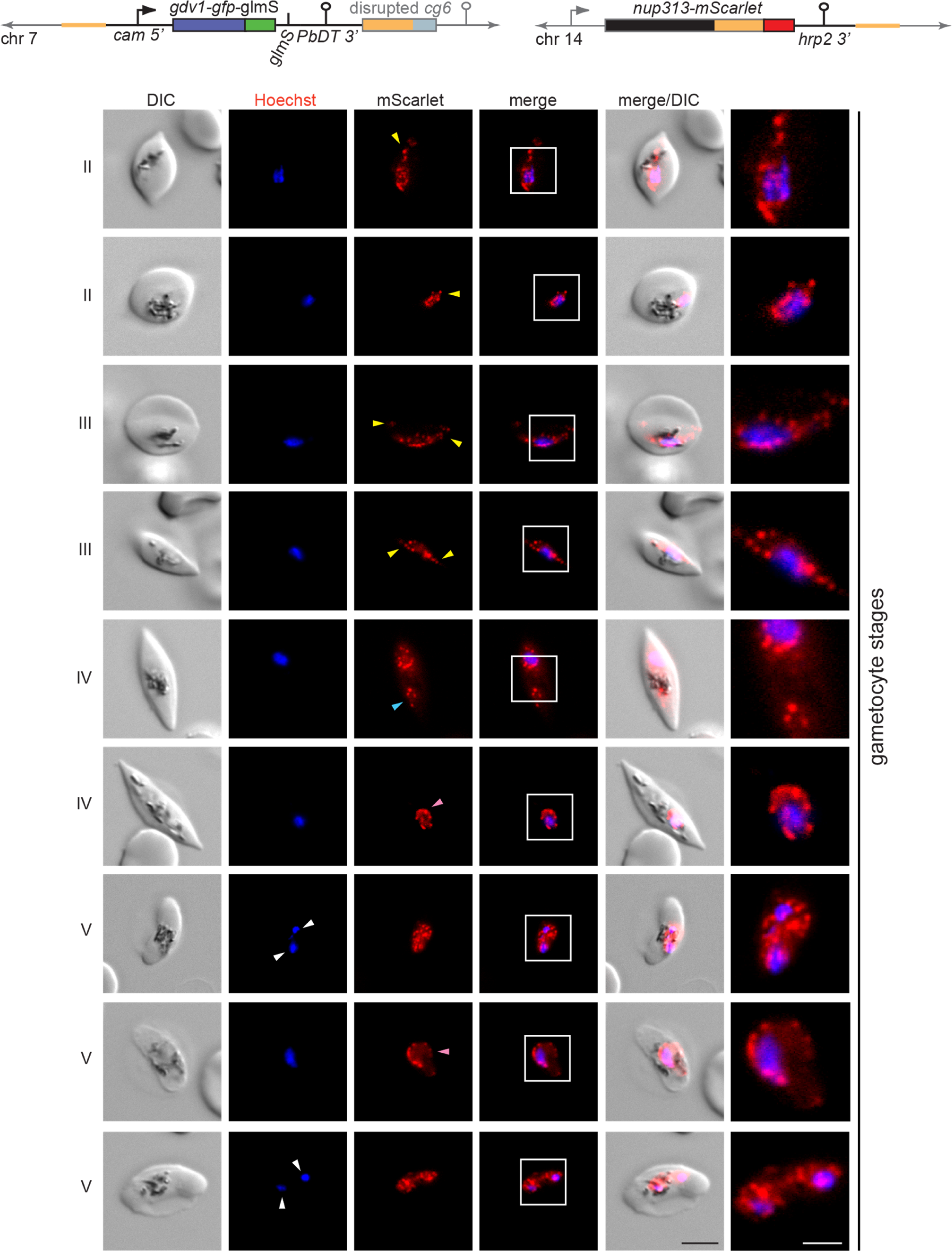
Nuclei in stage II to V gametocytes undergo marked morphological transformations. Schematic maps of the disrupted *cg6* (*glp3*) locus (PF3D7_0709200) (grey) carrying a single inducible GDV1-GFP-glmS expression cassette and the tagged *nup313* locus in double-transgenic NF54/iGP2_NUP313-mSc line are shown on top. The 5’ and 3’ homology regions used for CRISPR/Cas9-based gene editing are indicated in orange (see also Figs. S2 and S4). Representative live cell fluorescence microscopy images showing the localization of NUP313-mScarlet (red) in stage II to V gametocytes. Lateral extensions (yellow arrowheads) or rounded expansions (pink arrowheads) of the nucleus away from Hoechst-stained bulk chromatin and separate NUP313-mScarlet-delineated regions enclosing (white arrowheads) or not containing Hoechst-stained bulk chromatin (blue arrowhead) are highlighted. II-V, stage II to V gametocytes. DIC, differential interference contrast. Nuclei were stained with Hoechst (blue). Scale bar, 5 µm. White frames refer to the magnified view presented in the rightmost images (scale bar, 2 µm).

## Discussion

Gametocytes are the only forms of the malaria parasite that can secure human-to-human parasite transmission by the mosquito vector. While seminal research published in recent years provided major new insight into the regulatory mechanisms underlying sexual commitment and conversion ^5,26-31,33-35,37,38,65^, our molecular understanding of gametocyte biology is still unsatisfactory. New experimental tools conducive to the detailed investigation of gametocytes and their transmission to mosquitoes will be crucial to close these knowledge gaps and to inform and support the development of transmission-blocking drugs and vaccines in a best possible way.

Here, we engineered robust inducible gametocyte producer (iGP) lines that will allow investigating gametocyte biology with a similar level of routine, depth and detail hitherto reserved only for the study of asexual blood stage parasites. We achieved this by inserting a single conditional GDV1 overexpression cassette into the genomes of 3D7 and NF54 parasites. These iGP lines overcome the limitations of current protocols used for the bulk preparation of gametocytes. First, induction of sexual commitment is entirely independent of cumbersome and unreliable culture handling protocols and only requires adding Shield-1 and/or removing GlcN (depending on the specific iGP line used) from the culture medium for two days to induce GDV1 overexpression during the IDC preceding gametocyte differentiation. Second, the targeted induction of GDV1 overexpression triggers consistent sexual commitment rates of 75%, which allows generating large numbers of gametocytes even from small culture volumes. We cannot explain why GDV1 overexpression fails to trigger sexual commitment in the entire population but we assume GDV1 expression levels and/or the efficiency of GDV1-dependent HP1 eviction from the *pfap2-g* locus may vary between individual parasites. Third, sexual commitment is induced in a synchronous population of trophozoites, which leads to synchronous sexual differentiation in the progeny and therefore facilitates the preparation of stage-specific gametocyte populations across gametocyte maturation. Fourth, gametocyte synchronicity and yield are highly reproducible between experiments; starting with approximately 0.75×10^7^ iGP ring stage parasites in the commitment cycle (5% hematocrit, 1.5% parasitemia) routinely delivers up to 2×10^8^ stage V gametocytes per 10 ml of culture volume in a single induction experiment (Fig. 6), which increases gametocyte yields by at least one order of magnitude compared to previous reports ^58,63,86^. Because gametocyte production can be induced once every second day from an asexual feeder culture, the iGP lines developed here provide a constant rich source of synchronous gametocyte populations for experimental studies.

**Figure 6.**
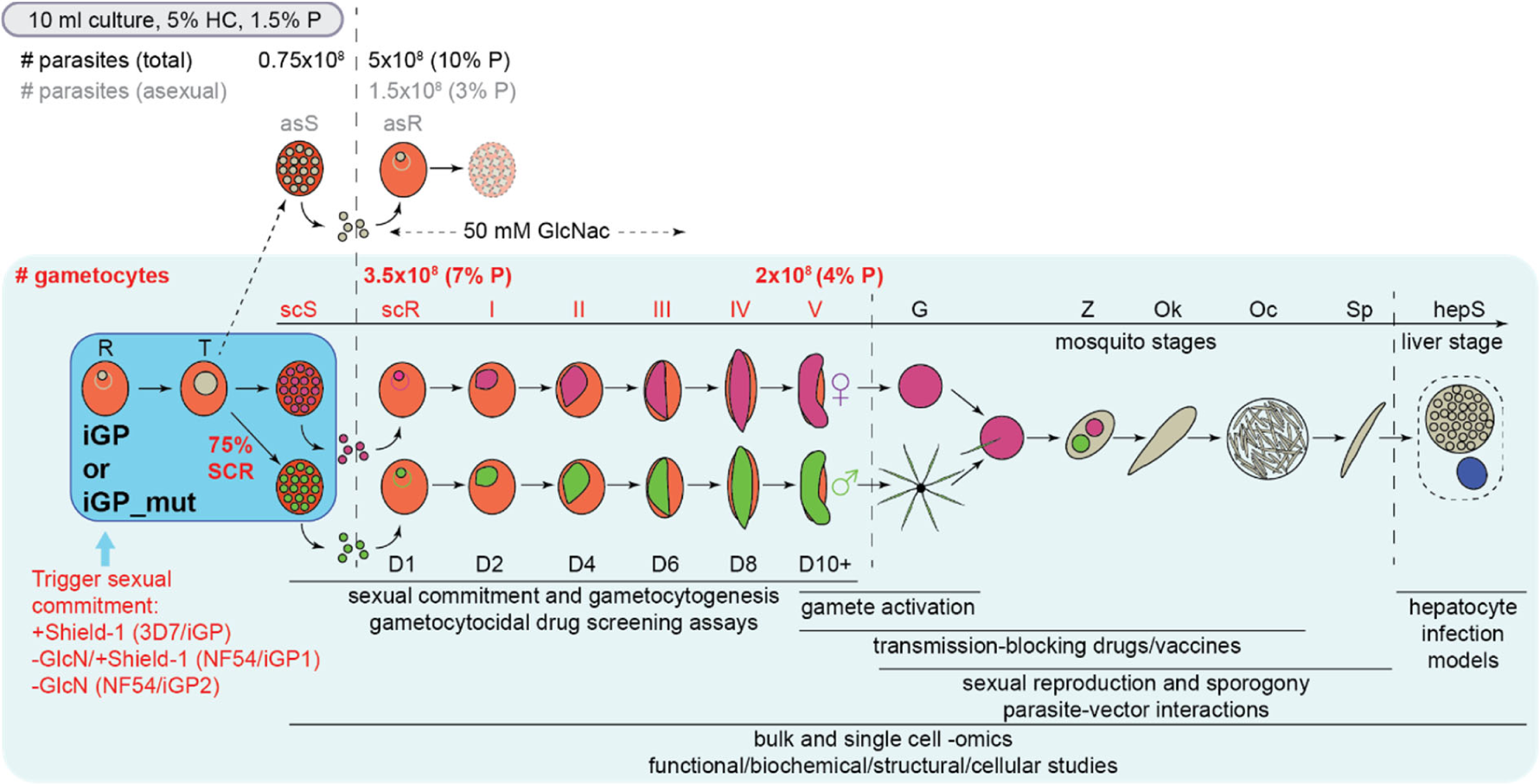
Scheme depicting the simple iGP induction protocol for the routine mass production of gametocytes and potential applications for future research. Addition of Shield-1 (3D7/iGP), addition of Shield-1/removal of GlcN (NF54/iGP1) or removal of GlcN (NF54/iGP2) from a synchronous ring stage culture triggers sexual commitment in trophozoites and produces ring stage progeny consisting of approximately 75% sexual ring stage parasites. Addition of 50 mM GlcNAc to the culture medium for the next six days eliminates remaining asexual parasites. Expected numbers of parasite-infected RBCs (#) and percent parasitaemia (% P) of total (black letters), asexual (grey letters) and sexual (red letters) parasites in the progeny routinely obtained from a 10 ml culture at 5% hematocrit (HC) and 1.5% starting parasitaemia are indicated. Asexual parasites are depicted in grey, sexually committed parasites and gametocytes are depicted in purple (females) and green (males). asS/scS, asexual/sexually committed schizont; asR/scR, asexual/sexually committed ring stage; T, trophozoite; I-V, gametocyte stages I-V; D1-D10, days 1-10 of gametocyte maturation; G, gametes; Z, zygote, Ok, ookinete; Oc, oocyst; Sp, sporozoites; hepS, intra-hepatic schizont. Possible applications of iGP lines for basic, applied and translational research on *P. falciparum* gametocytes and mosquito stage parasites are listed below the schematic.

The 3D7/iGP line produces exflagellation-defective male stage V gametocytes, similar to the recently engineered conditional gametocyte producer line E5ind that uses a conditional promoter swap approach to induce PfAP2-G expression ^37^. Both of these lines are therefore not useful to investigate the biology of late stage gametocytes and parasite stages in the mosquito or to conduct gametocyte transmission-blocking drug assays. However, they still provide excellent tools to dissect the molecular events linked to sexual conversion and early gametocyte differentiation. The E5ind line is superior in this regard as it displays virtually zero background sexual conversion and achieves 90% sexual conversion rates upon induction of PfAP2-G expression, which allows identifying molecular signatures of sexually committed parasites with high precision ^37^. The 3D7/iGP line will be useful to complement such studies and to dissect the mechanisms involved in GDV1-dependent activation of *pfap2-g*.

In contrast to the abovementioned cell lines, gametocytes obtained from the two NF54/iGP lines are infectious to *Anopheles* mosquitoes, which offers numerous additional opportunities for basic research and for the discovery and development of transmission-blocking interventions (Fig. 6). We placed great emphasis on constructing marker-free NF54/iGP lines and clones to allow further rounds of genetic manipulation using any of the three drug resistance markers routinely used for selection of transgenic parasites (h*dhfr, bsd*, yeast dihydroorotate dehydrogenase). This possibility provides greatest flexibility for the engineering of NF54/iGP mutants. Manipulating genes in NF54/iGP parasites offers the unprecedented advantage that the study of gene and protein function by phenotypic, cell biological, biochemical or structural analyses can routinely be performed not only in asexual blood stage parasites but readily also in a large number of isogenic gametocytes and possibly subsequent life cycle stages (Fig. 6). NF54/iGP lines and mutants will lend themselves for time-resolved high throughput profiling experiments to generate comprehensive transcriptomics, (phospho-)proteomics, epigenomics and metabolomics reference and experimental datasets for each stage of gametocyte development using both bulk and single-cell approaches to understand the regulatory processes governing male and female gametocyte differentiation. Furthermore, they will facilitate the targeted dissection of molecular mechanisms underlying the specific biology and metabolism of gametocytes. NF54/iGP lines will further be a suitable tool to investigate the differential susceptibility of immature and mature gametocytes to antimalarial drugs and will greatly simplify and streamline high throughput drug screening campaigns aiming to identify novel transmission-blocking compounds. Moreover, NF54/iGP gametocytes will support studies investigating sexual reproduction and subsequent parasite development in mosquitoes as well as parasite-vector interactions and vector competence. Potentially, NF54/iGP gametocytes may also be applicable for the development and validation of transmission-blocking vaccines and may provide a reliable resource for the production of sporozoites for hepatocyte infection models and for the optimization of protocols aiming to produce zygotes, ookinetes, oocysts and sporozoites *in vitro* ^87-89^ (Fig. 6). Noteworthy, the iGP approach is in principle also transferable to other *P. falciparum* strains through CRISPR/Cas9-mediated genomic insertion of an inducible GDV1 overexpression cassette. Generating iGP lines in different genetic backgrounds could facilitate the systematic analysis of potential strain-dependent differences in gametocyte biology, sensitivity to antimalarial drugs, infectiousness to mosquitoes or the capacity to undergo cross-fertilisation.

We illustrated the feasibility and benefit of subjecting NF54/iGP parasites to a second round of genetic engineering by tagging the nuclear pore component NUP313 in the NF54/iGP2 line. Our fluorescence microscopy data demonstrate marked differences in NPC abundance and distribution in the nuclear envelope as well as in nuclear morphology between asexual blood stage parasites and gametocytes (Figs. 4 and 5, Fig. S5). Our results on asexual parasites recapitulate previous findings obtained by high-resolution microscopy ^80,84,85^. These studies showed that merozoites and ring stage parasites possess only three to seven nuclear pores that are closely clustered in one region of the nuclear membrane. As ring stage parasites develop into trophozoites the number of NPCs increases and up to 60 NPCs are evenly distributed throughout the nuclear envelope. During schizogony the number of pores per nucleus decreases with increasing numbers of nuclei formed, such that in late schizonts each nucleus of the developing merozoites again possesses only a small number of clustered nuclear pores ^80,84,85^. The functional relevance of this intriguing redistribution of NPCs is currently unknown, but it may be linked to regulatory strategies of stage-specific gene expression during the IDC and/or the coordination of nuclear segregation during schizogony ^84^. In contrast, we found that gametocyte nuclei contain a stable and relatively high number of evenly distributed NPCs throughout all stages of gametocyte development, similar to what is observed for trophozoite nuclei. These observations are consistent with the open chromatin conformation and high transcriptional activity observed in both life cycle stages ^52,90,91^ and are suggestive of a high demand for nucleocytoplasmic shuttling during periods of growth. Importantly, because NPCs serve as a surrogate marker delineating the nuclear envelope our results also allow us to conclude that the nucleus in stage II to V gametocytes (i) undergoes marked morphological transformations reflected in narrow lateral extensions along the longitudinal axis of the gametocyte or rounded expansions of the nuclear envelope; and (ii) is frequently considerably larger than what might be extrapolated from the Hoechst-stained bulk of genetic material, in line with observations made in *P. berghei* gametocytes ^81^. In some late stage gametocytes we even observed two distinct NPC-demarcated regions, of which one or both stained positive with Hoechst. In trophozoites, such alterations in nuclear size and shape and spatial chromatin compaction has to our knowledge not been reported and was also not observed in this study. These data therefore suggest that nuclear reorganization and functional genome compartmentalization plays an important role in transcriptional reprogramming during gametocytogenesis and/or in the preparation for genome endoreplication and fertilisation in male and female stage V gametocytes, respectively. Interestingly, results obtained from TEM studies suggested that the nuclei of female and male gametocytes differ markedly both in size and shape, with the female nucleus being generally oval-shaped and comparably small and the male nucleus appearing substantially larger with lobular shape and distending towards the poles of the gametocyte ^12,54^. It is therefore tempting to speculate that the cells containing enlarged and irregularly shaped nuclei detected in our study may primarily represent male gametocytes. Unfortunately, we are unable to identify female and male gametocytes with confidence by live cell microscopy. Although differential haemozoin crystal distribution (dispersed in males, clustered in females) has been proposed as a diagnostic feature to distinguish between the two sexes ^7,12^, many gametocytes displayed ambiguous pigment patterns and we clearly observed irregular nuclear shapes in cells with clustered or dispersed haemozoin distribution (Figs. 4b and 5 and Fig. S5), suggesting that the nucleus undergoes marked transformations in both male and female gametocytes. Notwithstanding this uncertainty, our analysis of NF54/iGP2_NUP313-mSc parasites revealed first insight into the intriguing morphological features of gametocyte nuclei based on standard fluorescence microscopy and provides an excellent starting point for further detailed cytological and functional investigations of nuclear biology in *P. falciparum* transmission stages.

In summary, we engineered marker-free inducible gametocyte producer (iGP) lines that facilitate the routine mass production of synchronous gametocyte populations across all stages of gametocyte development using a most simple experimental setup and at a hitherto unprecedented scale. We demonstrate that NF54/iGP gametocytes are infectious to mosquitoes and that further genetic engineering of NF54/iGP parasites is straightforward. Owing to these favourable properties, we believe the iGP approach developed in this study will become an invaluable and broadly applicable tool for fundamental, applied and translational research on *P. falciparum* transmission stages.

## Materials and Methods

### Parasite culture

*P. falciparum* 3D7 parasites were cultured *in vitro* as described ^92^ using AB+ human RBCs at 5% haematocrit and RPMI 1640 medium supplemented with 25 mM HEPES, 100 mM hypoxanthine, 24 mM sodium bicarbonate and complemented with 0.5% Albumax II (Life Technologies). NF54 parasites were cultured with 10% AB+ human serum instead of 0.5% Albumax II. Growth medium was replaced daily. Intra-erythrocytic growth synchronization was achieved using repeated sorbitol treatments ^93^. The NF54/iGP1, NF54/iGP2 and NF54/iGP2_NUP-mScarlet lines were cultured in presence of 2.5 mM D-(+)-glucosamine hydrochloride (GlcN) to maintain *glmS* ribozyme activity during routine parasite propagation.

### Transfection constructs for CRISPR/Cas9-based gene editing

Inducible *gdv1* transgene expression cassettes were inserted into the *cg6* (*glp3*) locus (PF3D7_0709200) of 3D7 and NF54 wild type parasites by co-transfecting 50 μg each of a CRISPR/Cas9 transfection vector and a donor plasmid delivering the transgene cassette. 3D7/iGP1 was obtained by co-transfecting pHF_gC-*cg6* and pD_*cg6_cam-gdv1-gfp-dd* into 3D7 wild type parasites. NF54/iGP1 was obtained by co-transfecting pBF_gC-*cg6* and pD_*cg6_cam-gdv1-gfp-dd-glmS* into NF54 wild type parasites. NF54/iGP2 was obtained by co-transfecting pBF_gC-*cg6* and pD_*cg6_cam-gdv1-gfp-glmS* into NF54 wild type parasites. NF54/iGP2_NUP313mSc was obtained by co-transfecting pBF_gC-*nup313* and pD_*nup313-mScarlet* into NF54/iGP2 parasites. All transfection plasmids cloned in this study are derivatives of the original pHF_gC/pBF_gC CRISPR/Cas9 and pD donor plasmids published by Filarsky and colleagues ^34^ and are schematically displayed in Figs. S1, S2 and S4.

The pHF_gC-*cg6* CRISPR/Cas9 plasmid was generated by T4 DNA ligase-dependent insertion of annealed complementary oligonucleotides (11F, 11R) encoding the single guide RNA (sgRNA) target sequence sgt_*cg6* along with compatible single-stranded overhangs into *Bsa*I-digested pHF_gC ^34^. The sgt_*cg6* target sequence (gcacaaatataaattaaatt) is positioned 24-44 bp downstream of the start codon of the cg6 (*glp3)* gene and has been designed using CHOPCHOP ^94^. pBF_gC-*cg6* was generated in the same manner using the pBF_gC vector ^34^.

The pD_*cg6_cam-gdv1-gfp-dd* donor plasmid was generated by Gibson assembly ^95^ of five PCR fragments encoding (1) the *P. falciparum* calmodulin (*cam)* promoter followed by a *gdv1-gfp-dd* fusion gene (amplified from pHcam*-gdv1-gfp-dd* ^34^ using primers 1F and 1R); (2) the *P. berghei* dihydrofolate reductase-thymidylate synthase (*pbdhfr-ts*) terminator sequence (amplified from the pH_gC vector ^34^ using primers 2F and 2R); (3) a 387 bp *cg6* 3’ homology region (HR) spanning bps 392-778 of the *cg6* coding sequence (amplified from 3D7 gDNA using primers 3F and 3R); (4) the plasmid backbone (amplified from pUC19 using primers 4F and 4R); and (5) a 276 bp *cg6* 5’ HR spanning bps −343 to - 68 upstream of the *cg6* gene (amplified from 3D7 gDNA using primers 5F and 5R).

The pD_*cg6_cam-gdv1-gfp-dd-glmS* donor plasmid has been cloned by Gibson assembly using three fragments, namely (1) the *Eco*RI/*Age*I-fragment of pD_*cg6_cam-gdv1-gfp-dd* encoding part of the *cg6* 3’ HR (bps 482-778 of the *cg6* coding sequence), the vector backbone, the *cg6* 5’ HR, the *cam* promoter, the *gdv1* gene, and bps 1-715 of the *gfp* coding sequence; (2) a PCR fragment encoding bps 699-714 of the *gfp* coding sequence followed by the *dd* and *glmS-246* sequence (amplified from pD_*ap2g-gfp-dd-glmS* ^34^ using primers 7F and 7R); and (3) a PCR fragment encompassing the *pbdhfr-ts* terminator sequence and part of the *cg6* 3’ HR (spanning bps 392-501 of the *cg6* coding sequence) (amplified from pD_*cg6_cam-gdv1-gfp-dd* using primers 8F and 8R).

The pD_*cg6_cam-gdv1-gfp-glmS* vector was generated through Gibson assembly of two PCR fragments representing (1) the *pbdhfr-ts* terminator sequence, the *cg6* 3’ HR, the vector backbone, the *cg6* 5’ HR, the *cam* promoter, the *gdv1* gene, and bps 1-714 of the *gfp* coding sequence followed by a TAA stop codon (amplified from pD_*cg6_cam-gdv1-gfp-dd-glmS* using primers 9F and 9R); and (2) the *glmS-246* sequence (amplified from pD_*ap2g-gfp-dd-glmS* ^34^ using primers 10F and 10R).

To tag the nucleoporin NUP313 (PF3D7_1446500) with mScarlet ^96^ in NF54/iGP2 parasites, the following CRISPR/Cas9 and donor plasmids were cloned. The pBF_gC-*nup313* vector was generated by T4 DNA ligase-dependent insertion of annealed complementary oligonucleotides (18F, 18R) encoding the sgRNA target sequence sgt-*nup313* along with compatible single-stranded overhangs into *Bsa*I-digested pBF_gC ^34^. The sgt_*nup313* target sequence (gcactttgtagagataagta) is positioned at 91-110 bp downstream of the *nup313* coding sequence. The pD_*nup313-mScarlet* donor vector is a result of Gibson assembly of five PCR fragments encompassing (1) a 941 bp *nup313* 5’ HR spanning bps 8063-9003 of the *nup313* coding sequence (amplified from NF54 gDNA using primers 13F and 13R); (2) the *mScarlet* gene (amplified from a *P. falciparum* codon-adjusted synthetic sequence (Genscript) (Fig. S6) using primers 14F and 14R); (3) the *P. falciparum* histidine-rich protein 2 (*hrp2)* terminator sequence (amplified from pBF_gC with primers 15F and 15R); (4) a 1’000 bp *nup313* 3’ HR spanning the region 90-1089 bp downstream of the *nup313* coding sequence (amplified from NF54 gDNA using primers 16F and 16R); and (5) the vector backbone amplified from pBF_gC using primers 17F and 17R. All oligonucleotide sequences used for cloning and Sanger sequencing are provided in Table S1.

### Parasite transfection and selection of transgenic lines

Parasites were transfected and gene-edited parasites selected as described ^34^. The NF54/iGP1 and NF54/iGP2 transgenic lines were cultivated in presence of 2.5 mM GlcN to maintain activity of *glmS* ribozyme in the *gdv1-gfp-dd-glmS* and *gdv1-gfp*-*glmS* transgene cassettes. Drug selection started 24 hours post transfection. 3D7/iGP parasites were selected with 5 nM WR99210 (WR) for the first six days and NF54/iGP1, NF54/iGP2 and NF54/iGP2_NUP313-mSc parasites were selected with 2.5 µg/ml blasticidin-S-HCl (BSD) for the first eleven days. Thereafter, all transfected parasites were cultured in absence of drug pressure until stably propagating populations were established. PCR on gDNA was performed to confirm proper genome editing and absence of plasmid DNA. The 3D7/iGP and NF54/iGP1 lines, which still retained the pHF_gC or pBF_gC CRISPR/Cas9 plasmids expressing the negative selection marker yFCU fused to hDHFR or BSD deaminase, respectively, were treated with 40 µM 5-fluorocytosine (5-FC) to obtain plasmid-free populations. 3D7/iGP, NF54/iGP1 and NF54/iGP2 lines were cloned out by limiting dilution using a plaque assays as described ^97^. All oligonucleotide sequences used for PCR on parasite gDNA and plasmid DNA are provided in Table S1.

### Fluorescence microscopy

α-GDV1-GFP-DD and α-Pfs16 IFAs were performed with methanol-fixed cells using mouse mAbs α-GFP (1:200) (Roche, #11814460001) and α-Pfs16 (1:500) ^98^ primary antibodies, and Alexa Fluor 488 goat α-mouse IgG secondary antibody (1:200) (Invitrogen, **#**A-11001). Staining of DNA and mounting of IFA slides was performed using Vectashield with DAPI (Vector Laboratories, #H-1200). Microscopy was performed using a Leica DM 5000B microscope with a Leica DFC 345 FX camera using a 63x immersion oil objective (total magnification = 1008x). All images were acquired via the Leica Application Suite software (version LAS 4.9.0) and processed using Adobe Photoshop CC with identical settings.

To perform live cell fluorescence imaging of NUP313-mScarlet presented in Fig. 4 and Fig. S5, asexual blood stages were stained with 1 µg/mL DAPI (Biomol) in RPMI and gametocytes were stained with 4.5 µg/mL Hoechst33342 (Chemodex) in PBS at 37°C for 15 min. Stained cells were pelleted at 1000 g for 1 min and the pellet resuspended in an equal volume of supernatant. 10 µL of the sample were placed on a microscopy slide and covered with a cover slip. Fluorescence microscopy images were acquired with a Leica DM6 B microscope equipped with Leica DFC9000 GT camera using a 100x immersion oil objective (total magnification = 1000x). Filter block settings: mScarlet (Ex. 542-585nm; Em. 604-644nm), DAPI/Hoechst (Ex. 325-375; Em. 435-485). All images were acquired with the Leica Application Suite X software (LAS X) and processed using Adobe Photoshop CS2 with identical settings.

To perform live cell fluorescence imaging of NUP313-mScarlet presented in Fig. 5, gametocytes were stained with 4.5 µg/mL Hoechst33342 (Sigma-Aldrich) in PBS at 37 °C for 20 min. Stained cells were pelleted at 300 g for 1 min and the pellet resuspended in an equal volume of the supernatant. 3 µL of the sample were placed on a microscopy slide, mixed with 3 µL Vectashield and covered with a cover slip. Images were acquired with a Leica DM 5000B microscope and a Leica DFC 345 FX camera using a 100x immersion oil objective (total magnification = 1250x). Filter block settings: mScarlet (Ex. 543/22 nm; Em. 593/40 nm), Hoechst (Ex. 377/50 nm; Em. 447/60). All images were acquired using the Leica Application Suite software (LAS 4.9.0) and processed using ImageJ (version 1.52n) using identical settings.

### Induction of sexual commitment and gametocytogenesis

Depending on the inducible GDV1 overexpression system employed, induction of sexual commitment was achieved by either stabilizing GDV1-GFP-DD fusion protein expression via addition of Shield-1 (3D7/iGP), stabilization of *gdv1-gfp* mRNA via removal of GlcN from the culture medium (NF54/iGP2 and NF54/iGP2_NUP313-mSc), or employing both approaches simultaneously (NF54/iGP1). To determine sexual conversion rates, synchronous ring stage cultures at 1-3 % parasitaemia (0-16 hpi, generation 1) were washed in culture medium and split into two identical cultures at 5% haematocrit. One was maintained under non-inducing conditions (control population representing background sexual conversion rates) and the other was induced for sexual conversion by triggering ectopic GDV1-GFP(-DD) expression through addition of either 1350, 675 or 337.5 nM Shield-1 (3D7/iGP), addition of 1350 nM Shield-1/removal of GlcN (NF54/iGP1), or removal of GlcN (NF54/iGP2, NF54/iGP2_NUP313-mSc). The inducing conditions were reversed 54 hrs later in the ring stage progeny (6-22 hpi, generation 2; day 1 of gametocytogenesis). To estimate sexual conversion rates parasites were methanol-fixed at 36-44 hpi in generation 2 (day 2 of gametocytogenesis) and α-Pfs16 IFAs combined with DAPI staining were performed. Conversion rates were determined based on fluorescence microscopy counts of early stage I gametocytes (Pfs16- and DAPI-positive) compared to all infected RBCs (DAPI-positive).

For gametocyte maturation assays, parasites were induced for sexual conversion as describe above. Culture medium containing 50 mM N-acetyl-D-glucosamine (GlcNAc) was added to the ring stage progeny (generation 2; day 1 of gametocytogenesis) and changed daily for six days to eliminate asexual parasites ^58,72^. From day seven onwards gametocytes were cultured with normal culture medium. Gametocyte stages and gametocytaemia were assessed by visual inspection of Giemsa-stained thin blood smears prepared until day 12 of gametocytogenesis using a 100x immersion oil objective (total magnification = 1000x).

### Standard Membrane Feeding Assays (SMFAs)

On day 10, 13 and 14 of gametocytogenesis aliquots of stage V gametocyte cultures were tested in the Standard Membrane Feeding Assay (SMFA) ^73^ according to an established workflow ^75^. Briefly, after a final change of culture medium on the day of mosquito feeding a small volume of the gametocyte culture was used to prepare the blood meal and to make blood smears. 300 µL of the gametocyte culture (approximately 3% gametocytaemia, 5% haematocrit) was mixed with pre-warmed 180 µL packed RBCs and the cells were pelleted for 20 sec. The supernatant was carefully removed and the RBC pellet resuspended in 150 µL pre-warmed human serum. Small membrane feeders were used to feed female *Anopheles stephensi* mosquitoes separately with each of the prepared gametocyte populations (approximately 0.3% gametocytaemia, 55% haematocrit). After eight days, midguts of 20 mosquitoes per feed were dissected and stained using 3% mercurochrome. The number of oocysts per midgut were counted by microscopy (1000x magnification).

## Data availability

All data generated or analysed during this study are included in this published article and its Supplementary Information files.

## Acknowledgements

This work was supported by the Swiss National Science Foundation (grant numbers BSCGI0_157729 and 31003A_169347) and the Fondation Pasteur Suisse. A.A. received a scholarship from the Jürgen Manchot Foundation.

## Author contribution

S.D.B. cloned all transfection plasmids, generated and characterised all transgenic cell lines and designed, performed and analysed experiments related to the NF54/iGP lines. A.P. designed, performed and analysed experiments related to the 3D7/iGP line. A.A and N.M.B.B performed fluorescence microscopy on the NF54/iGP2_NUP313-mSc line. M.V.B. performed and analysed mosquito infection experiments. T.W.G, H.P.B and R.W.S. provided conceptual advice and resources and supervised experiments. S.D.B., A.P., A.A. and N.M.B.B. helped preparing illustrations. T.S.V. conceived the study, designed, supervised, and analysed experiments, provided resources, prepared illustrations and wrote the paper. All authors contributed to editing of the manuscript.

