## Supplementary Information for "CRISPR/Cas9-engineered inducible gametocyte producer lines as a novel tool for basic and applied research on *Plasmodium falciparum* malaria transmission stages"

**This file includes:**

- Supplementary figures S1 to S6
- Supplementary table S1

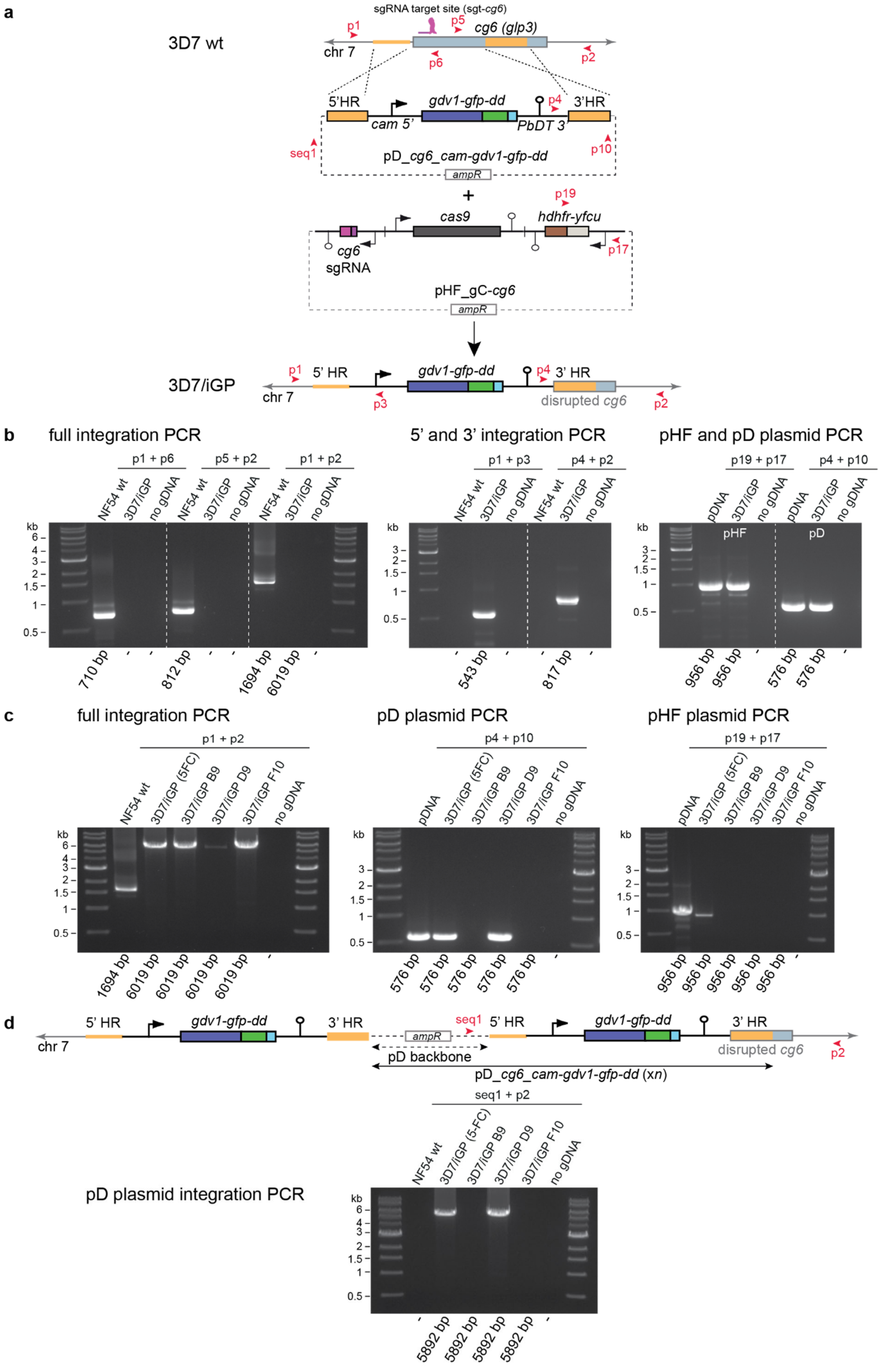
a
3D7 wt
sgRNA target site (sgt-cg6)
p1
p5
cg6 (glp3)
chr 7
p6
p2
5'HR
gdv1-gfp-dd
p4
3'HR
cam 5'
PbDT 3'
seq1
p10
pD_cg6_cam-gdv1-gfp-dd
ampR
+
p19
cas9
hdhfr-yfcu
cg6
sgRNA
p17
pHF_gC-cg6
ampR
3D7/iGP
p1
5' HR
gdv1-gfp-dd
p4
3' HR
chr 7
p3
disrupted cg6
p2
b
full integration PCR
p1 + p6
p5 + p2
p1 + p2
NF54 wt
3D7/iGP
no gDNA
710 bp
812 bp
1694 bp
6019 bp
5' and 3' integration PCR
p1 + p3
p4 + p2
543 bp
817 bp
pHF and pD plasmid PCR
p19 + p17
p4 + p10
pDNA
pHF
pD
956 bp
576 bp
kb
6
4
3
2
1.5
1
0.5
c
full integration PCR
p1 + p2
NF54 wt
3D7/iGP (5FC)
3D7/iGP B9
3D7/iGP D9
3D7/iGP F10
no gDNA
1694 bp
6019 bp
pD plasmid PCR
p4 + p10
576 bp
pHF plasmid PCR
p19 + p17
956 bp
d
5' HR
gdv1-gfp-dd
3' HR
ampR
seq1
chr 7
pD backbone
disrupted cg6
p2
pD_cg6_cam-gdv1-gfp-dd (xn)
seq1 + p2
3D7/iGP (5-FC)
pD plasmid integration PCR
5892 bp

**Supplementary Figure 1. CRISPR/Cas9-based engineering of the 3D7/iGP line. a** Schematic maps of the endogenous *cg6* (*glp3*) locus (PF3D7_0709200) in 3D7 wild type (wt) parasites (top), the pD_*cg6_cam-gdv1-gfp-dd* donor and pHF_gC-*cg6* CRISPR/Cas9 transfection plasmids (center), and the disrupted *cg6* locus carrying the inducible GDV1-GFP-DD expression cassette in 3D7/iGP parasites (bottom). The relative position of the sgt_*cg6* sgRNA target sequence is shown in purple. The pD_*cg6_cam-gdv1-gfp-dd* donor plasmid contains the *gdv1-gfp-dd* fusion gene controlled by the *P. falciparum cam* promoter and *pbdhfr-ts* terminator (PbDT 3') elements, flanked on either side by a homology region (HR) for homology-directed repair (orange). The pHF_gC-*cg6* plasmid contains expression cassettes for SpCas9 (dark grey), the sgRNA (purple) and the *hdhfr-fcu* positive-negative drug selection marker (brown-grey). Primer binding sites used to confirm successful gene editing by PCR are indicated by red arrowheads. **b** Diagnostic PCRs on gDNA from NF54 wt parasites and the 3D7/iGP mother line. Primer combinations to detect the presence and absence of the *cg6* wt locus in NF54 wt and 3D7/iGP, respectively, and integration of the full *gdv1-gfp-dd* expression cassette in 3D7/iGP (left panel). Primer combinations to detect the 5' and 3' recombination events in 3D7/iGP (middle panel). Primer combinations to detect the presence of the pHF_gC-*cg6* and the pD_*cg6_cam-gdv1-gfp-dd* plasmids in 3D7/iGP (right panel). pDNA, plasmid DNA control. **c** Diagnostic PCRs on gDNA from NF54 wt parasites, the 5-FC-treated 3D7/iGP mother line and three clones. Primer combinations to detect the presence and absence of the *cg6* wt locus in NF54 wt and 3D7/iGP parasites, respectively, and integration of the full *gdv1-gfp-dd* expression cassette in 3D7/iGP parasites (left panel). Primer combinations to detect the presence of the pD_*cg6_cam-gdv1-gfp-dd* (middle panel) and pHF_gC-*cg6* (right panel) plasmids in 3D7/iGP parasites. pDNA, plasmid DNA control. **d** Schematic map of a pD_*cg6_cam-gdv1-gfp-dd* donor plasmid concatamer integrated into the *cg6* locus by double-crossover recombination (integration of a tandem *gdv1-gfp-dd* expression cassette is shown). The *E. coli* plasmid backbone is indicated by a dashed arrow. The binding sites of the seq1 and p2 primers used to detect this event by PCR are indicated by red arrowheads. Diagnostic PCRs on gDNA from NF54 wt parasites, the 5-FC-treated 3D7/iGP mother line and three clones show the presence of an integrated pD_*cg6_cam-gdv1-gfp-dd* donor plasmid concatamer in the 5-FC-treated 3D7/iGP mother line and clone D9 but not in clones B9 and F10.

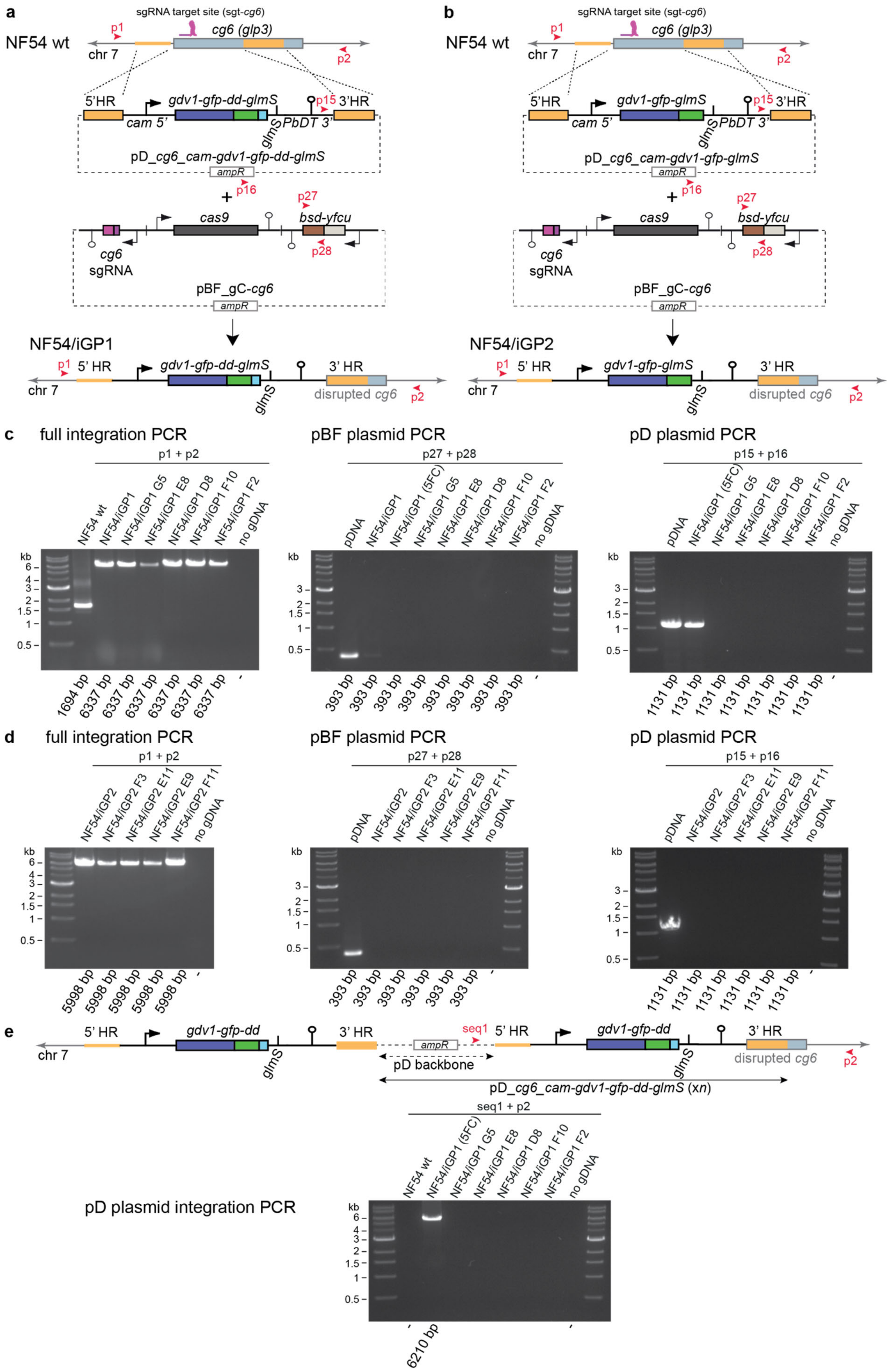
a
sgRNA target site (sgt-cg6)
p1
cg6 (glp3)
NF54 wt
chr 7
p2
5'HR
gdv1-gfp-dd-glmS
p15
3'HR
cam 5'
glmS
PbDT 3'
pD_cg6_cam-gdv1-gfp-dd-glmS
ampR
+
p16
p27
cas9
bsd-yfcu
cg6
sgRNA
p28
pBF_gC-cg6
ampR
NF54/iGP1
p1
5' HR
gdv1-gfp-dd-glmS
3' HR
chr 7
glmS
disrupted cg6
p2
b
sgRNA target site (sgt-cg6)
p1
cg6 (glp3)
NF54 wt
chr 7
p2
5'HR
gdv1-gfp-glmS
p15
3'HR
cam 5'
glmS
PbDT 3'
pD_cg6_cam-gdv1-gfp-glmS
ampR
+
p16
p27
cas9
bsd-yfcu
cg6
sgRNA
p28
pBF_gC-cg6
ampR
NF54/iGP2
p1
5' HR
gdv1-gfp-glmS
3' HR
chr 7
glmS
disrupted cg6
p2
c
full integration PCR
p1 + p2
NF54 wt
NF54/iGP1
NF54/iGP1 G5
NF54/iGP1 E8
NF54/iGP1 D8
NF54/iGP1 F10
NF54/iGP1 F2
no gDNA
kb
6
4
3
2
1.5
1
0.5
1694 bp
6337 bp
6337 bp
6337 bp
6337 bp
6337 bp
6337 bp
-
pBF plasmid PCR
p27 + p28
pDNA
NF54/iGP1
NF54/iGP1 (5FC)
NF54/iGP1 G5
NF54/iGP1 E8
NF54/iGP1 D8
NF54/iGP1 F10
NF54/iGP1 F2
no gDNA
kb
3
2
1.5
1
0.5
393 bp
393 bp
393 bp
393 bp
393 bp
393 bp
393 bp
393 bp
-
pD plasmid PCR
p15 + p16
pDNA
NF54/iGP1 (5FC)
NF54/iGP1 G5
NF54/iGP1 E8
NF54/iGP1 D8
NF54/iGP1 F10
NF54/iGP1 F2
no gDNA
kb
3
2
1.5
1
0.5
1131 bp
1131 bp
1131 bp
1131 bp
1131 bp
1131 bp
1131 bp
-
d
full integration PCR
p1 + p2
NF54/iGP2
NF54/iGP2 F3
NF54/iGP2 E11
NF54/iGP2 E9
NF54/iGP2 F11
no gDNA
kb
6
4
3
2
1.5
1
0.5
5998 bp
5998 bp
5998 bp
5998 bp
5998 bp
-
pBF plasmid PCR
p27 + p28
pDNA
NF54/iGP2
NF54/iGP2 F3
NF54/iGP2 E11
NF54/iGP2 E9
NF54/iGP2 F11
no gDNA
kb
3
2
1.5
1
0.5
393 bp
393 bp
393 bp
393 bp
393 bp
393 bp
-
pD plasmid PCR
p15 + p16
pDNA
NF54/iGP2
NF54/iGP2 F3
NF54/iGP2 E11
NF54/iGP2 E9
NF54/iGP2 F11
no gDNA
kb
3
2
1.5
1
0.5
1131 bp
1131 bp
1131 bp
1131 bp
1131 bp
1131 bp
-
e
5' HR
gdv1-gfp-dd
3' HR
ampR
seq1
5' HR
gdv1-gfp-dd
3' HR
chr 7
glmS
pD backbone
glmS
disrupted cg6
p2
pD_cg6_cam-gdv1-gfp-dd-glmS (xn)
seq1 + p2
NF54 wt
NF54/iGP1 (5FC)
NF54/iGP1 G5
NF54/iGP1 E8
NF54/iGP1 D8
NF54/iGP1 F10
NF54/iGP1 F2
no gDNA
pD plasmid integration PCR
kb
6
4
3
2
1.5
1
0.5
-
6210 bp
-

**Supplementary Figure 2. CRISPR/Cas9-based engineering of the NF54/iGP lines. a** Schematic maps of the endogenous *cg6* (*glp3*) locus (PF3D7_0709200) in NF54 wild type (wt) parasites (top), the pD_*cg6_cam-gdv1-gfp-dd-glmS* donor and pBF_gC-*cg6* CRISPR/Cas9 transfection plasmids (center), and the disrupted *cg6* locus carrying the inducible GDV1-GFP-DD-glmS expression cassette in NF54/iGP1 parasites (bottom). The relative position of the sgt_*cg6* sgRNA target sequence is shown in purple. The pD_*cg6_cam-gdv1-gfp-dd-glmS* donor plasmid contains the *gdv1-gfp-dd-glmS* fusion gene controlled by the *P. falciparum cam* promoter and *P. berghei dhfr-ts* terminator (PbDT 3'), flanked on either side by a homology region (HR) for homology-directed repair (orange). The pBF_gC-*cg6* plasmid contains expression cassettes for SpCas9 (dark grey), the sgRNA (purple) and the *bsd-fcu* positive-negative drug selection marker (brown-grey). Primer binding sites used to confirm successful gene editing by PCR are indicated by red arrowheads. **b** Schematic maps of the endogenous *cg6* (*glp3*) locus (PF3D7_0709200) in NF54 wt parasites (top), the pD_*cg6_cam-gdv1-gfp-glmS* donor and pBF_gC-*cg6* CRISPR/Cas9 transfection plasmids (center), and the disrupted *cg6* locus carrying the inducible GDV1-GFP-glmS expression cassette in NF54/iGP2 parasites (bottom). **c** Diagnostic PCRs on gDNA from NF54 wt parasites, the NF54/iGP1 mother line before and after 5-FC treatment and five NF54/iGP1 clones. Primer combinations to detect the presence and absence of the *cg6* wt locus in NF54 wt and NF54/iGP1 parasites, respectively, and integration of the full *gdv1-gfp-dd* expression cassette in NF54/iGP1 parasites (left panel). Primer combinations to detect the presence of the pBF_gC-*cg6* (middle panel) and pD_*cg6_cam-gdv1-gfp-dd-glmS* (right panel) plasmids. **d** Diagnostic PCRs on gDNA from the NF54/iGP2 mother line and four NF54/iGP2 clones. Primer combinations are as described above for panel c. **e** Schematic map of a pD_*cg6_cam-gdv1-gfp-dd-glmS* donor plasmid concatamer integrated into the *cg6* locus by double-crossover recombination (integration of a tandem *gdv1-gfp-dd-glmS* expression cassette is shown). The *E. coli* plasmid backbone is indicated by a dashed arrow. The binding sites of the seq1 and p2 primers used to detect this event by PCR are indicated by red arrowheads. Diagnostic PCRs on gDNA from NF54 wt parasites, the 5-FC-treated NF54//iGP1 mother line and five clones show the presence of an integrated pD_*cg6_cam-gdv1-gfp-dd-glmS* donor plasmid concatamer in the 5-FC-treated NF54/iGP1 mother line but not in any of the clones.

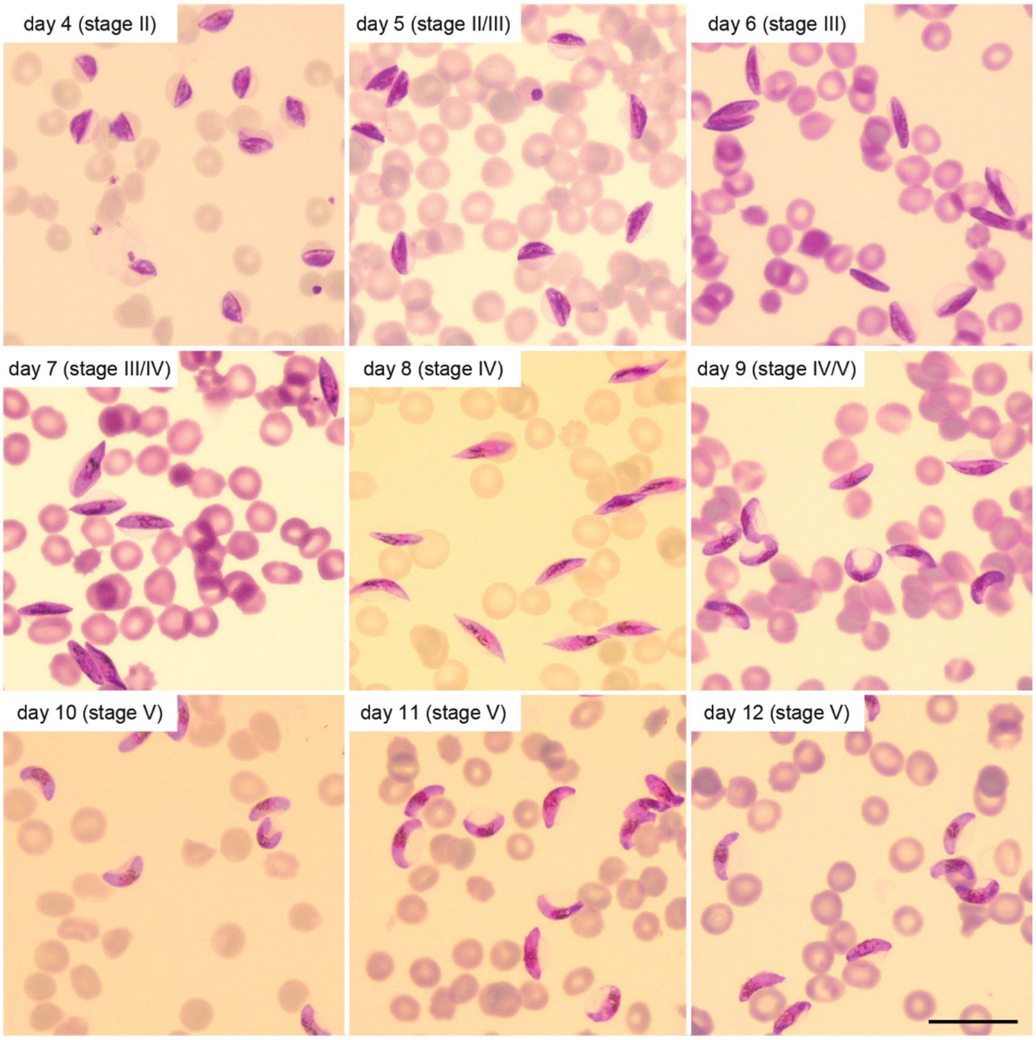


**Supplementary Figure 3. Synchronous maturation of induced NF54/iGP2 gametocytes.** Representative images of synchronously developing NF54/iGP2 gametocytes obtained after inducing sexual commitment in the previous cell cycle through removal of GlcN from the culture medium. Giemsa-stained blood smears were prepared daily from day 4 (stage II) to day 12 (stage V) of gametocytogenesis. Gametocyte cultures were treated with 50 mM GlcNAc from day 1 to 6 of gametocytogenesis to eliminate asexual blood stage parasites. Scale bar, 20 µm.

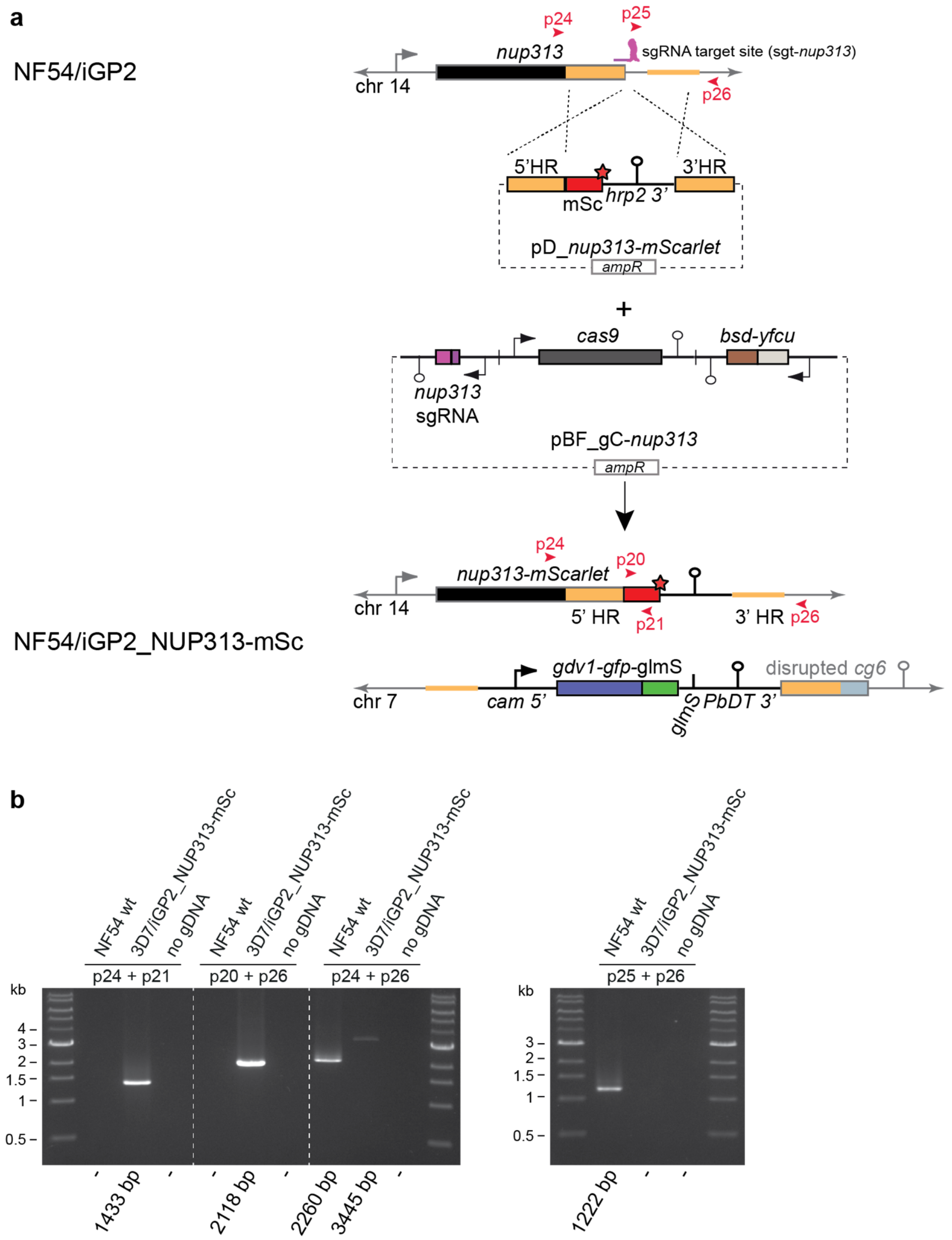


**Supplementary Figure 4. CRISPR/Cas9-based engineering of the NF54/iGP2_NUP313-mSc line.** **a** Schematic maps of the endogenous *nup313* locus (PF3D7_1446500) in NF54/iGP2 parasites (top), the pD_*nup313-mScarlet* donor and pBF_gC-*nup313* CRISPR/Cas9 transfection plasmids (center), and the edited *nup313* locus in NF54/iGP2_NUP313-mSc parasites expressing the NUP313-mScarlet fusion protein (bottom). The relative position of the sgt_*cg6* sgRNA target sequence is shown in purple. The pD_*nup313-mScarlet* donor plasmid contains the *mScarlet* sequence followed by the *P. falciparum* histidine-rich protein 2 (*hrp2*) terminator, flanked on either side by a homology region (HR) for homology-directed repair (orange). The 5' HR corresponds to the 3' end of *nup313* (omitting the stop codon), fused in frame to the *mScarlet* sequence. The pBF_gC-*nup313* plasmid contains expression cassettes for SpCas9 (dark grey), the sgRNA (purple) and the *bsd-fcu* positive-negative drug selection marker (brown-grey). Primer binding sites used to confirm successful gene editing by PCR are indicated by red arrowheads. The red asterisk denotes the STOP codon. **b** Diagnostic PCRs on gDNA from NF54 wild type (wt) and NF54/iGP2_NUP313-mSc parasites. Primer combinations to detect the 5' and 3' recombination events in NF54/iGP2_NUP313-mSc and the presence and absence of the *nup313* wt locus in NF54 wt and NF54/iGP2_NUP313-mSc, respectively, and insertion of the *mScarlet-hrp2 3'* sequence in NF54/iGP2_NUP313-mSc (left panel). Alternative primer combination to detect the presence and absence of the *nup313* wt locus in NF54 wt and NF54/iGP2_NUP313-mSc, respectively (right panel).

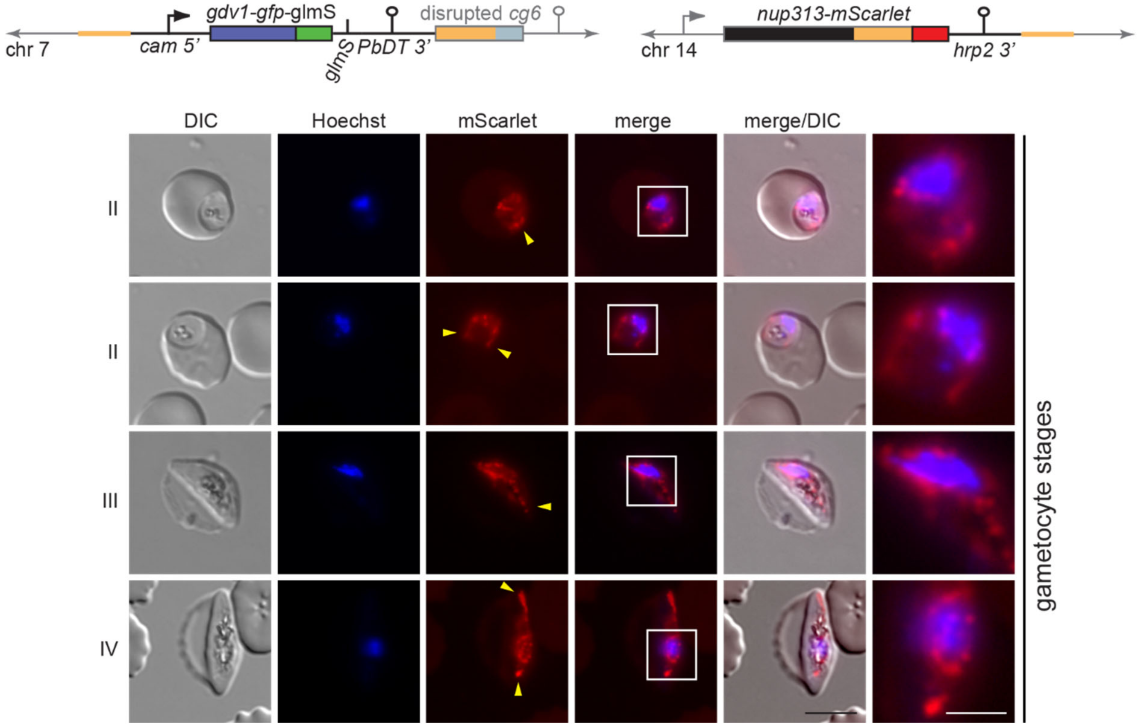


**Supplementary Figure 5. Nuclei in stage II to IV gametocytes undergo marked morphological transformations.** Schematic maps of the disrupted *cg6* (*glp3*) locus (PF3D7_0709200) (grey) carrying a single inducible GDV1-GFP-glmS expression cassette and the tagged *nup313* locus in double-transgenic NF54/iGP2_NUP313-mSc line are shown on top. The 5' and 3' homology regions used for CRISPR/Cas9-based gene editing are indicated in orange (see also Figs. S2 and S4). Representative live cell fluorescence microscopy images showing the localization of NUP313-mScarlet (red) in stage II to IV gametocytes. Lateral extensions of the nucleus away from Hoechst-stained bulk chromatin are highlighted by yellow arrowheads. II-IV, stage II to IV gametocytes. DIC, differential interference contrast. Nuclei were stained with Hoechst (blue). Scale bar, 5 µm. White frames refer to the magnified view presented in the rightmost images (scale bar, 2 µm).

**>mScarlet**
atggtgagcaagggcgaggcagtgatcaaggagttcatgcggttcaaggtgcacatggagggctccatgaacggccacgagttcgagatcgagg
gcgagggcgagggccgcccctacgagggcacccagaccgccaagctgaaggtgaccaagggtggcccctgcccttctcctgggacatcctgtc
ccctcagttcatgtacggctccagggccttcaccaagcaccccgccgacatccccgactactataagcagtccttccccgagggcttcaagtgg
gagcgcgtgatgaacttcgaggacggcggcgccgtgaccgtgacccaggacacctccctggaggacggcaccctgatctacaaggtgaagctcc
gcggcaccaacttccctcctgacggccccgtaatgcagaagaagacaatgggctgggaagcgtccaccgagcggttgtaccccgaggacggcgt
gctgaagggcgacattaagatggccctgcgcctgaaggacggcggccgctacctggcggacttcaagaccacctacaaggccaagaagcccgtg
cagatgcccggcgcctacaacgtcgaccgcaagttggacatcacctcccacaacgaggactacaccgtggtggaacagtacgaacgctccgagg
gccgccactccaccggcggcatggacgagctgtacaag

**>mScarlet co**
agtaaaggtgaagcagttataaaagaatttatgagatttaaagtacatatggaaggttcaatgaatggacatgaatttgaaatagaaggagaag
gtgaaggaagaccatatgaaggaacacaaacagctaaattgaaagttacaaaaggtggaccattaccatttagttgggatattttatcaccaca
atttatgtatggtagtagagcatttacaaaacatccagctgatataccagattattataaacaatcatttccagaaggatttaaatgggaaaga
gtaatgaattttgaagatggaggtgcagttacagtaacacaagatacaagtttagaagatggtacattaatttataaagttaaattgagaggta
caaattttccaccagatggaccagtaatgcaaaagaaaacaatgggatgggaagcatcaacagaaagattatatccagaagatggtgttttaaa
aggagatataaaaatggctttaagattaaaagatggaggtagatatttagcagattttaaaacaacatataaagctaaaaaaccagttcaaatg
ccaggtgcttataatgtagatagaaaattggatataacaagtcataatgaagattatacagttgtagaacaatatgaaagaagtgaaggaagac
attcaacaggtggaatggatgaattatataaa

**>Pairwise sequence alignment**
```
mScarlet      atggtgagcaagggcgaggcagtgatcaaggagttcatgcggttcaaggtgcacatggag      60
mScarlet co   ------agtaaaggtgaagcagttataaaagaatttatgagatttaaagtacatatggaa      54
                    ** ** ** ** ***** ** ** ** ** *** * ** ** ** ** *****

mScarlet      ggctccatgaacggccacgagttcgagatcgagggcgagggcgagggccgcccctacgag      120
mScarlet co   ggttcaatgaatggacatgaatttgaaatagaaggagaaggtgaaggaagaccatatgaa      114
              ** ** ***** ** ** ** ** ** ** ** ** ** ** ** **  * ** ** **

mScarlet      ggcacccagaccgccaagctgaaggtgaccaagggtggcccctgcccttctcctgggac      180
mScarlet co   ggaacacaaacagctaaattgaaagttacaaaaggtggaccattaccatttagttgggat      174
              ** ** ** ** ** **  **** ** ** ** ***** **  * ** **    *****

mScarlet      atcctgtcccctcagttcatgtacggctccagggccttcaccaagcaccccgccgacatc      240
mScarlet co   attttatcaccacaatttatgtatggtagtagagcatttacaaaacatccagctgatata      234
              **  * ** ** ** ** ***** **    ** ** ** ** ** ** ** ** ** **

mScarlet      cccgactactataagcagtccttccccgagggcttcaagtgggagcgcgtgatgaacttc      300
mScarlet co   ccagattattataaacaatcatttccagaaggatttaaatgggaaagagtaatgaatttt      294
              ** ** ** ***** ** ** ** ** ** ** ** ** *****  * ** ***** **

mScarlet      gaggacggcggcgccgtgaccgtgacccaggacacctccctggaggacggcaccctgatc      360
mScarlet co   gaagatggaggtgcagttacagtaacacaagatacaagtttagaagatggtacattaatt      354
              ** ** ** ** ** ** ** ** ** ** ** **     * ** ** ** **  * **

mScarlet      tacaaggtgaagctccgcggcaccaacttccctcctgacggccccgtaatgcagaagaag      420
mScarlet co   tataaagttaaattgagaggtacaaattttccaccagatggaccagtaatgcaaaagaaa      414
              ** ** ** **  *  * ** ** ** ** ** ** ** ** ** ******** *****

mScarlet      acaatgggctgggaagcgtccaccgagcggttgtaccccgaggacggcgtgctgaagggc      480
mScarlet co   acaatgggatgggaagcatcaacagaaagattatatccagaagatggtgttttaaaagga      474
              ******** ******** ** ** **  * ** ** ** ** ** ** **  * ** **

mScarlet      gacattaagatggccctgcgcctgaaggacggcggccgctacctggcggacttcaagacc      540
mScarlet co   gatataaaaatggctttaagattaaaagatggaggtagatatttagcagattttaaaaca      534
              ** ** ** *****  *  *  * ** ** ** **  * **  * ** ** ** ** **

mScarlet      acctacaaggccaagaagcccgtgcagatgcccggcgcctacaacgtcgaccgcaagttg      600
mScarlet co   acatataaagctaaaaaaccagttcaaatgccaggtgcttataatgtagatagaaaattg      594
              ** ** ** ** ** ** ** ** ** ***** ** ** ** ** ** **  * ** ***

mScarlet      gacatcacctcccacaacgaggactacaccgtggtggaacagtacgaacgctccgagggc      660
mScarlet co   gatataacaagtcataatgaagattatacagttgtagaacaatatgaaagaagtgaagga      654
              ** ** **    ** ** ** ** ** ** ** ** ***** ** *** *    ** **

mScarlet      cgccactccaccggcggcatggacgagctgtacaag                              696
mScarlet co   agacattcaacaggtggaatggatgaattatataaa                              690
               * ** ** ** ** ***** **  * ** **
```

**Supplementary Figure 6. Pairwise alignment of the *mScarlet* and *P. falciparum* codon-optimised *mScarlet* gene sequences.** Nucleotide sequences and Clustal Omega (https://www.ebi.ac.uk/Tools/msa/clustalo/) output of the pairwise sequence alignment of *mScarlet* and the *P. falciparum* codon-optimised version of *mScarlet* (mScarlet co). Asterisks denote identical nucleotides.

### Supplementary Table 1. Oligonucleotides used in this study.

| Application | Primer name | Target sequence | Sequence (5'-3') |
|---|---|---|---|
| PCRs to clone transfection vectors | 1F | *cam* promoter | gaataaataaatataaatatgaacatgatcttataaggaaattccc |
| | 1R | *dd* cds | gaacattaagctgccatatcctcattccagttttagaagc |
| | 2F | *pbdhfr-ts* terminator | gagcttctaaaactggaatgaggatatggcagcttaatg |
| | 2R | *pbdhfr-ts* terminator | tcatgctaccctgaagaagaaaag |
| | 3F | *cg6* cds | cttcttcagggtagcatgaacatgtaaaagataaaaatg |
| | 3R | *cg6* cds | cctcttcgctattacgccagcaacttgtctatgccacc |
| | 4F | plasmid backbone | ctggcgtaatagcgaagagg |
| | 4R | plasmid backbone | cattaatgaatcggccaacg |
| | 5F | *cg6* upstream | gggaatttccttataagatcatgttcatatttatatttatttattcatttgtttg |
| | 5R | *cg6* upstream | cgttggccgattcattaatgcacatattcgctccttc |
| | 7F | *gfp/dd cds* | ggatgaactatacaaaaccggttctagtatgggagtgcaggtggaaac |
| | 7R | *glmS* | cgaacattaagctgccatatccgctagcatttttcttcctcc |
| | 8F | *pbdhfr-ts* terminator | aggaggaagaaaaatgctagcggatatggcagcttaatgttcg |
| | 8R | *cg6* cds | ctcttctactctttcgaattcaccatgttcatcttttatacatttatc |
| | 9F | *glmS/pbdhfr-ts* terminator | aggaggaagaaaaatgctagcggatatggcagcttaatgttcg |
| | 9R | *gfp* cds | ctattgagaaaataagaacaagattatttgtatagttcatccatgccatg |
| | 10F | *glmS* sequence | gcatggatgaactatacaaataatcttgttcttattttctcaatag |
| | 10R | *glmS* sequence | cgaacattaagctgccatatccgctagcatttttcttcctcctaagattg |
| | 13F | *nup313* cds | agtgagcgaggaagcggaagctgtttctaattcaagatctgattcc |
| | 13R | *nup313* cds | acttgtggatccaccactactatttatcatattttgattcataaatttatgcc |
| | 14F | *mScarlet* cds | atagtagtggtggatccacaagtaaaggagaagcagttataaaag |
| | 14R | *mScarlet* cds | tctattattaaataaatttatttgtatagttcatccattccacc |
| | 15F | *pfhrpII* terminator | ggaatggatgaactatacaaataaatttatttaataatagattaaaaatattataaaaataaaaac |
| | 15R | *pfhrpII* terminator | gcactttgtagagataagtaatttaataaatatgttcttatatataatgag |
| | 16F | *nup313* downstream | gaacatatttattaaattacttatctctacaaagtgcacacac |
| | 16R | *nup313* downstream | gcaccatatgcgaaggagaaatttcctttgaggac |
| | 17F | plasmid backbone | gtcctcaaaggaaatttctccttcgcatatggtgcactctcagtac |
| | 17R | plasmid backbone | gaatcagatcttgaattagaaacagcttccgcttcctcgctcac |
| sgRNA annealing | 11F | sgt_*cg6* | tattgcacaaatataaattaaatt |
| | 11R | sgt_*cg6* | aaacaatttaatttatatttgtgc |
| | 18F | *nup313 downstream* | aaactacttatctctacaaagtgc |
| | 18R | *nup313 downstream* | tattgcactttgtagagataagta |
| PCRs to verify gene editing | p1 | *cg6* upstream | atgtagcccatgaaagagttatg |
| | p2 | *cg6* downstream | cacaagcacataatggtggg |
| | p3 | *cam* promoter (only in pD) | gcatgcaagcttcgatcc |
| | p4 | *pbdhfr-ts* terminator | gctcaattctttatgtccacaac |
| | p5 | *cg6* cds | ggtagagttcaattcatcaaacc |
| | p6 | *cg6* cds | gatcctgggtaacttcacag |
| | p10 | pHF/pBF/pD backbone | gtactgagagtgcaccatatgc |
| | p15 | *pbdhfr-ts* terminator | gatattgagcagaggatatgc |
| | p16 | pHF/pBF/pD backbone | gcacccaactgatcttcagc |
| | p17 | *cam* promoter (only in pHF/pBF) | gctcgcaaatggccaaataag |
| | p19 | *hdhfr* cds | ccttgtggaggttccttgag |
| | p20 | *mScarlet* cds | ggaggtgcagttacagtaacacaag |
| | p21 | *mScarlet* cds | gcattactggtccatctggtgga |
| | p24 | *nup313* cds | tgagcatatagtaccatcagaatgg |
| | p25 | *nup313* downstream | acacaataaaatgttcacggaataatg |
| | p26 | *nup313* downstream (Pf3D7_1446600 cds) | gatacaagggaaggaatacaacg |
| | p27 | *bsd* cds | atggcacctttgtctcaagaag |
| | p28 | *bsd* cds | accctcccacacataaccag |
| | seq 1 | pD backbone (except pD_*nup313-mScarlet*) | gcgaggaagcggaagagc |
